# When shared concept cells support associations: theory of overlapping memory engrams

**DOI:** 10.1101/2021.03.12.434964

**Authors:** Chiara Gastaldi, Tilo Schwalger, Emanuela De Falco, Rodrigo Quian Quiroga, Wulfram Gerstner

**Author notes:** TS and WG jointly supervised this work. communicating author (Chiara Gastaldi).

## Abstract

Assemblies of neurons, called concepts cells, encode acquired concepts in human Medial Temporal Lobe. Those concept cells that are shared between two assemblies have been hypothesized to encode associations between concepts. Here we test this hypothesis in a computational model of attractor neural networks. We find that for concepts encoded in sparse neural assemblies there is a minimal fraction *c*_min_ of neurons shared between assemblies below which associations cannot be reliably implemented; and a maximal fraction *c*_max_ of shared neurons above which single concepts can no longer be retrieved. In the presence of a periodically modulated background signal, such as hippocampal oscillations, recall takes the form of association chains reminiscent of those postulated by theories of free recall of words. Predictions of an iterative overlap-generating model match experimental data on the number of concepts to which a neuron responds.

**Authors contributions:** All authors contributed to conception of the study and writing of the manuscript. CG and TS developed the theory. CG wrote the code for all figures. EDF and RQQ provided the experimental data. EDF and CG analyzed the data. WG and CG developed algorithms to fit the experimental data.

## Introduction

Human memory exploits associations between concepts. If you visited a famous place with a friend, a postcard of that place will remind you of him or her. The episode “with my friend at this place” has given rise to an association between two existing concepts: before the trip (the episodic event), you already knew your friend (first concept) and had seen the place (second concept), but only after the trip, you associate these two concepts.

Concepts are encoded in the human Medium Temporal Lobe (MTL) by neurons, called “concept cells”, that respond selectively and invariantly to stimuli representing a specific person or a specific place [1–3]. Each concept is thought to be represented by an assembly of concept cells that increases their firing rates simultaneously upon presentation of an appropriate stimulus. The fraction *γ* of neurons in the human MTL which is involved in the representation of each concept is estimated to be *γ* ∼0.23% [4]. Under the assumption that each memory item is represented by the activation of a fixed, but random, subset of active neurons, a single concept is expected to activate *γN* neurons and two arbitrary concepts are expected to share *γ*^2^*N* cells, where *N* is the total number of neurons in the relevant brain areas.

Experimental studies have shown that single neurons can become responsive to new concepts while learning pairs of associations [5]. Moreover, it has been estimated that assemblies representing two arbitrary concepts share less than 1% of neurons, whereas assemblies representing previously associated concepts share about 4−5% of neurons [6] suggesting that an increased fraction of shared neurons supports the association between concepts [6–8].

With the presence of shared neurons, the activation of a first assembly (e.g., a place) may also activate a second assembly (e.g., a person). This poses several theoretical questions. First, for the brain to function correctly as a memory network, it must remain possible to recall the two associated concepts separately (e.g. place without your friend), and not automatically the two together. However, if the concepts share too many neurons it becomes likely that the two memory items can no longer be distinguished, but are merged into a single, broader concept encoded by a larger number of active neurons. We therefore ask as a first question: what is the maximally allowed fraction *c*_max_ of shared neurons between two assemblies before the possibility of separate memory recalls breaks down? Shared concept cells can be visualised as an overlap between two memory engrams. Below the maximal fraction *c*_max_ of shared neurons, each of the associated patterns can be recalled as a separate memory pattern, as schematically illustrated in Fig. 1A.

**Figure 1.**
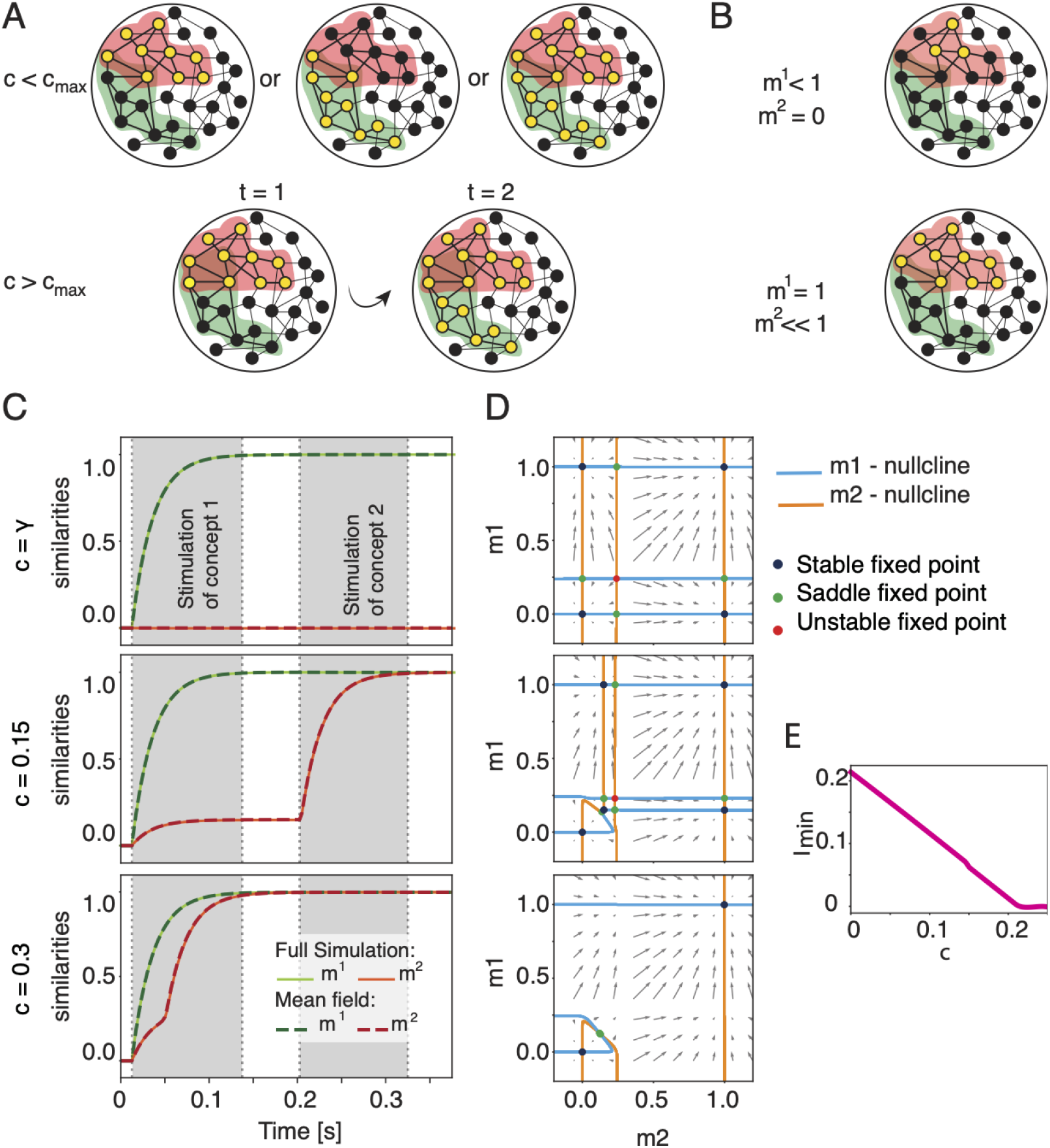
Overlapping concepts can be retrieved separately and jointly. A) Activation of concepts (schematic). Black filled circles = inactive neurons. Yellow filled circles = active neurons. Colored halos (red, green) represents assignment to a specific concept. When the fraction of shared neurons is small (top row, *c < c*_max_) the two concepts can be recalled separately or together. If the number of shared concept cells is too large (bottom row, *c > c*_max_), the recall of a first concept (red) leads inevitably to the activation of the second associated concept (green). B) Similarity measure. If only a subset of neurons belonging to the first memory engram is activated (top), the configuration exhibits similarities *m*^1^ *<* 1 and *m*^2^ = 0. If the first memory is fully recalled, while memory 2 is not (bottom), the similarity measures are *m*^1^ = 1 and *m*^2^ *<<* 1. C) Dynamics of the similarities for different fractions of shared neurons. The similarities *m*^1^ (green) and *m*^2^ (red) as a function of time in a full network simulation (solid lines) are compared to predictions of mean-field theory (dashed lines). Strong external stimulation *I*_1_ = 0.3 is given to the units belonging to concept *µ* = 1 during a first stimulation period and a weak external stimulation *I*_2_ = 0.1 is given to the units belonging to concept *µ* = 2 during the second stimulation period (in grey). If *c > c*_max_, the concept 2 gets activated without receiving any stimulation. D) Three phase-planes of the dynamics of similarity variables *m*^1^ and *m*^2^ for different values of fraction of shared neurons *c*. Arrows indicate direction and speed of increase or decrease of the similarity variables. Intersections of blue and orange lines (the “nullclines” of the two variables *m*^1^, *m*^2^) indicate fixed points, with a stability encoded by color (legend). E) Minimum amplitude of the external stimulation *I*^2^ needed to activate the memory of the second concept if the first one is activated (as a function of the fraction of shared neurons*c*). Parameters: 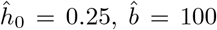, *r*_max_ = 40 Hz, *τ* = 25 ms, *α* = 0, *γ* = 0.2%. For simulations: *N* = 10000, *P* = 2.

As an alternative to a *static* recall of one or the other concept (or the two associated concepts together), we could also ask whether the activation of a concept would facilitate the recall of an associated one, or even a *temporal* chain activation of associations (as described in free memory recall tasks [9–12]), due to overlaps in the representations. In this context, we ask a second question: if each concept is represented by a small fraction of active neurons *γ*, given the activation of a concept, is there a minimal fraction of shared neurons *c*_min_ necessary to enable a reliable activation of associated ones?

Moreover, while most experimental studies have dealt with pairwise associations between, say one person and one place, more recent work has shown that a single neuron can respond to multiple concepts [6], e.g., several related places. In view of this, we ask a third question: how should memory be organized in a neural network such that *k* different memory engrams all have the equal size pairwise overlaps?

Associative memory in recurrent networks, such as the area CA3 of the hippocampus, has been modeled with attractor neural networks [13–17] where each memory item is encoded as a memory engram [18, 19] in a fixed random subset of neurons (called “pattern” in the theoretical literature [17]) such that no pattern has an overlap above chance with another one. Animal studies provide evidence of attractor dynamics in area CA3 [20, 21]. The few theoretical studies that considered overlapping memory engrams above chance level in the past [22, 23] focused on overlaps arising from a hierarchical organization of memories. Whereas such a hierarchical approach is suitable for modeling memory representation in the cortex, we are interested in modeling MTL, and in particular area CA3 of the hippocampus, where experimentally no hierarchical or topographical organization has been observed [6]. In experiments, episodic associations between *arbitrary* different concepts (such as a person and a place) - and shared neurons in the corresponding assemblies - can be induced by joint presentation of images representing the different concepts [5]. Inspired by these experiments, we create pairwise associations between a number of concepts by artificially introducing shared concept cells in the model. We will talk about “overlapping engrams” if the number of shared concept cells is beyond the number *γ*^2^*N* of cells that are shared by chance.

## Results

The first two questions introduced above can be summarized as a more general one: What is the role of those concept cells that are shared between stored memory engrams? To answer this question, we consider an attractor neural network of *N* neurons in which *P* engrams are stored in the form of binary random patterns [7]. The pattern 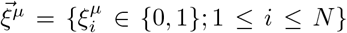 with pattern index *µ* ∈ {1, …, *P*} represents one of the stored memory engrams: a value 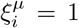 indicates that neuron *i* is part of the stored memory engram and therefore belongs to the assembly of concept *µ*, while a value of 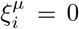 indicates that it does not. A network that has stored *P* memory engrams is said to have a memory load of *α* = *P/N*.

Since concept-cells in human hippocampus form sparse neural assemblies with a sparseness parameter *γ* ∼ 0.23% [4], we focus on the case of sparse memory engrams. In other words, an arbitrary neuron *i* has a low probability 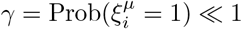 to participate in the assembly of concept cells corresponding to memory engram *µ*.

The attractor neural network is implemented in a standard way [24, 25]. Each neuron, *i* = 1, …, *N*, is modelled by a firing rate model [25]

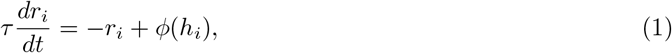

where *r*_*i*_(*t*) is the firing rate of neuron *i* and *φ*(*h*) = *r*_max_*/*{1 + exp[−*b*(*h* − *h*_0_)]} is the sigmoidal transfer function, or frequency-current (f-I) curve, characterized by the firing threshold *h*_0_, the maximal steepness *b*, and the maximal firing rate *r*_max_. The patterns 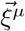 are encoded in the synaptic weights *w*_*ij*_ via a Hopfield-Tsodyks connectivity for sparse patterns so that the average of synaptic weights across a large population of neurons vanishes [17].

In attractor neural network models, memory engrams *µ* induce stable values 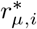 of the neuronal firing rates during the retrieval of a stored concept. In mathematical terms, to each engram *µ* corresponds a fixed point 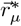 in such a way that the firing rate 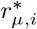 of neuron *i* is high if 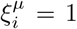 and low if 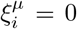. When the network state 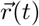 is initialized close enough to the stored memory *µ*, the attractor dynamics drives the network to the retrieval state 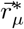 characterized by persistent activity of all those neurons that belong to the assembly of concept *µ*.

The similarity between the momentary network state and a stored memory *µ* is defined as

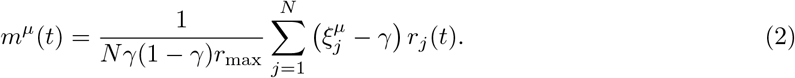

The similarity measures the correlation between the firing rates {*r*_*j*_(*t*)}_*j* = 1,…,*N*_ and the stored patterns 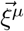 such that if memory concept *µ* is retrieved, then *m*^*µ*^∼ 1 (schematics in Fig. 1B), and, if no memory is recalled (*resting state*), then *m*^*µ*^∼ 0 for all *µ*. The similarity of the network activity with a stored memory develops as a function of time. For example, computer simulations of a network of *N* = 10 000 interacting neurons indicate that, if one of two engrams that share concept cells is stimulated for 120ms, then the similarity of the network activity with this engram increases to a value close to one, indicating that the memory has been recalled (Fig. 1C middle) while the second memory is only weakly activated quantified by a small, but non-zero similarity. However, if the fraction of shared neurons is above a maximally allowed fraction *c*_max_, then the second memory always gets activated even before it is stimulated (Fig. 1C bottom) indicating that associations are so strong that the two concepts have been merged.

### Maximal fraction of shared neurons between memory engrams

In order to better understand the network dynamics, we develop a mathematical theory that depends on the fraction of neurons *c* that are shared between two engrams. The *total number n of shared neurons* in a network of size *N* depends on *c* and the sparsity parameter *γ*, i.e., *n* = *γcN*.

Let us imagine to gradually increase the fraction of shared neurons between the first two memory engrams. At the lowest end, *c* = *γ*, the patterns 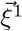 and 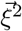 are independent, and hence cell assemblies 1 and 2 share a small fraction of neurons corresponding to chance level. It is well known, that in this case, each memory engram generates a separate attractive fixed-point of the network dynamics [17] implying that the two corresponding concepts can be retrieved separately. However, experimental data reports that, for associated concepts, the fraction of shared neurons *c*∼4−5% [6] is much larger than the chance level *γ*∼0.23%. This observation suggests that the patterns 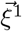 and 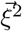 of two associated memory engrams have a fraction of shared neurons larger than chance level, *c > γ*. On the other hand, in the (trivial) limit case of large fraction of shared neurons *c* → 1, the two memory engrams and hence the two cell assemblies share all neurons, and it is clearly impossible to retrieve one memory without the other.

To study the maximal fraction of shared neurons *c*_max_ at which independent memory recall breaks down, we use a mean-field approach for large networks and work in the limit *N*→ ∞. In this limit, it is possible to fully describe the network dynamics using the similarities *m*^*µ*^ as the relevant macroscopic variables. Since we are interested in the retrieval process of concepts *µ* =1 and 2, we assume the similarity of the present network state with other memories *µ >* 2 to be close to zero: we will refer to these non-activated memories as “background patterns”. Under these assumptions, we find dynamical mean-field equations that capture the network dynamics through the similarity variables *m*^1^ and *m*^2^.

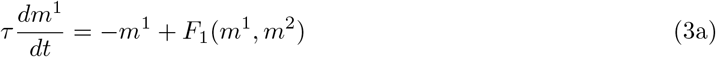

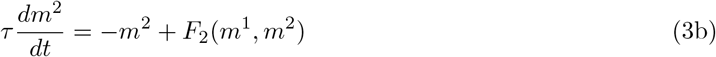

where the explicit form of the functions *F*_1_ and *F*_2_ is given in Eq. (10) of Methods. Equation (3) represents a two-dimensional dynamical systems which can be analyzed using phase-plane analysis. Figure 1D shows three phase-planes in the *m*^1^−*m*^2^ space, each for a different value of the fraction of shared neurons. The *m*^1^- or *m*^2^-nullclines solve *dm*^1^*/dt* = 0 or *dm*^2^*/dt* = 0 in Eq. (3a) and (3b), respectively. The intersections between the *m*^1^- and *m*^2^-nullcline are equilibrium solutions, or fixed points, of the mean-field dynamics and are color-coded according to their stability. For *c* = *γ*, we identify four stable fixed points: the resting state (*m*^1^, *m*^2^) = (0, 0), two single-retrieval states (*m*^1^, *m*^2^) = (1, 0) and (*m*^1^, *m*^2^) = (0, 1) corresponding to the retrieval of concept *µ* = 1 and the retrieval of concept *µ* = 2, respectively. Finally, there is a symmetric state which corresponds to the activation of both concepts simultaneously, (*m*^1^ = *m*^2^ ≲ 1).

Once a maximally allowed value *c* = *c*_max_ is reached, the two single-retrieval states merge with their nearby saddle points and disappear. To compute the numerical value of the maximal fraction of shared neurons, we extract it from the bifurcation diagram (Fig. S1). For fractions of shared neurons *c > c*_max_ only two stable fixed points are left, the resting state and the symmetric state in which assemblies of both concepts are activated together: this symmetric state is the theoretical description of the state that we qualitatively predicted above where the activation of a first concept leads inevitably to the activation of the second, overlapping one (Fig. 1C, bottom). The minimum external stimulation needed to activate the second concept depends on the fraction of shared neurons (Fig. 1E). With our choice of parameters, no external stimulation is needed to recall the second memory, if the fraction of shared neurons is *c > c*_max_ = 22%, since the two concepts have merged into a single one and are *always* recalled together.

In the limit of infinite steepness *b* → ∞, vanishing load *α* = 0 and vanishing sparseness *γ* → 0, the value 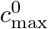 of the maximal fraction of shared neurons can be calculated analytically. Since this value provides an upper bound of the maximal fraction of shared neurons for arbitrary *b*, we have the inequality (Fig. 2A)

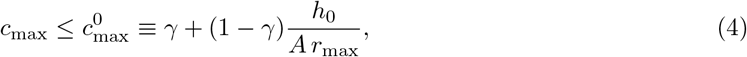

where *A* characterizes the overall strength of synaptic weights (see Eq.(5) below). Further analysis (see Methods) shows that the stationary states of the mean-field dynamics depend – apart from the parameters *γ, C* and *α* related to the patterns – only on two dimensionless parameters: the rescaled firing threshold *ĥ* _0_ = *h*_0_*/*(*A r*_max_) and the rescaled steepness 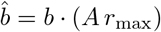. We find that the maximal fraction of shared neurons *c*_max_ increases with 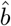 and, for *ĥ* _0_ *<* 0.8, also with *ĥ* _0_ (Fig.2A).

**Figure 2.**
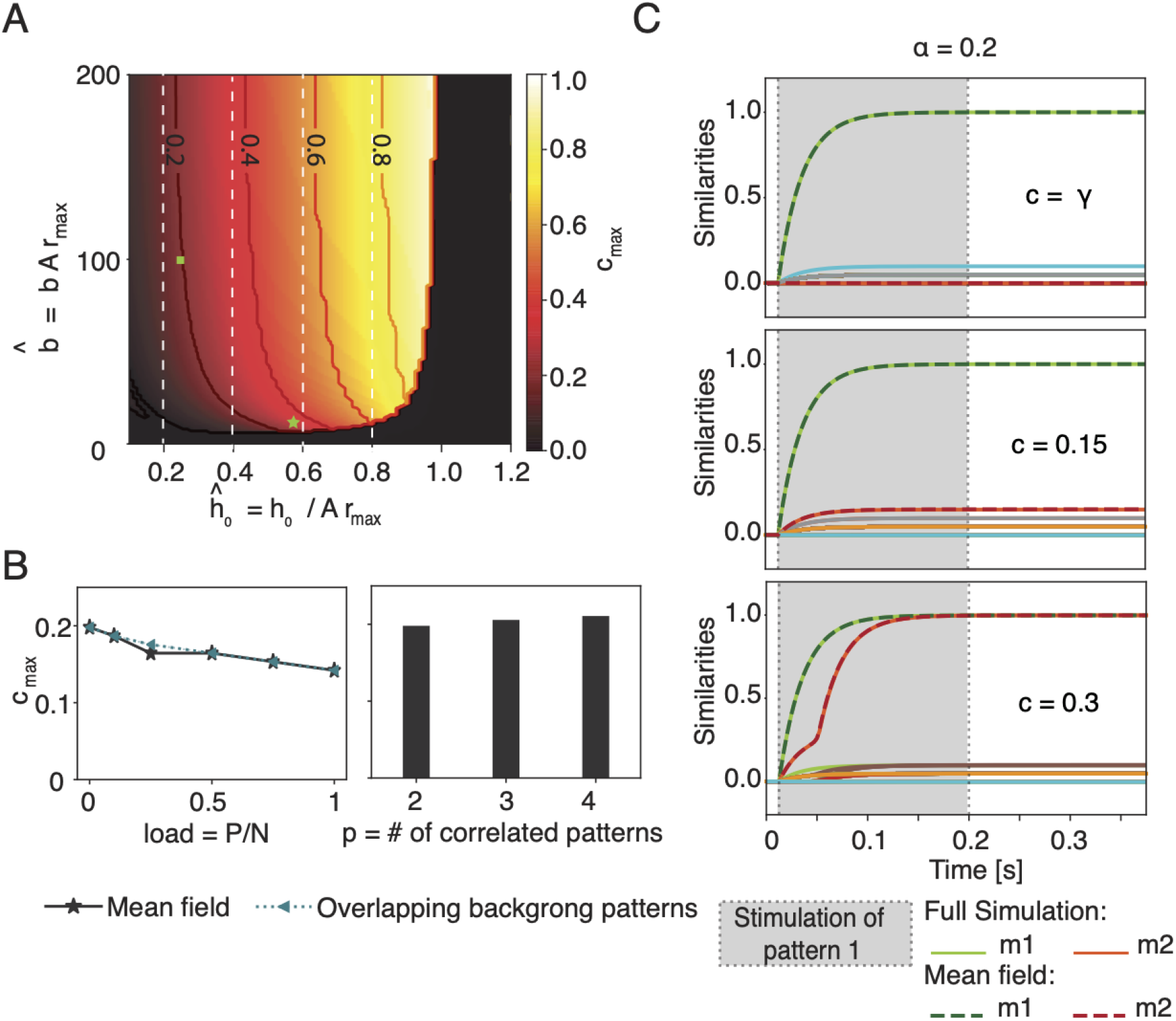
The maximal fraction *c*_max_ of shared neurons depends on the neuronal frequency-current curve but not on the memory load. A) Maximal fraction (*c*_max_, color code) as a function of the parameters 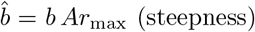 and *ĥ* _0_ = *h*_0_*/*(*Ar*_max_) (firing threshold) of the frequency-current curve. Niveau lines added for indicated values of *c*_max_. In the black area the resting state is the only stable solution.Vertical white dashed lines indicate the theoretical upper bound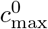, for different values of *ĥ*_0_. The green square indicates the parameter choice used in Fig. 1 and 2B-C. The green star indicates the parameters extracted for the Macaque inferotemporal cortex [24]. B) Maximal fraction *c*_max_ of shared neurons as a function of the memory load *α* = *P/N* (left graph) without (solid grey line) or with overlaps in pairs of two of the *P*−2 background patters (dashed green line); and as a function of the number *p* of correlated patterns (histogram, right graph). C) As in Fig. 1C, but with a large number of background patterns (*α* = 0.2). Network activity exhibits only small similarity with background patterns (diversely colored lines) but large similarity with the stimulated pattern *µ* = 1. Parameters (unless specified): 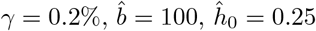, *r*_max_ = 40 Hz, *τ* = 25ms; *α* = 0 in A-B. For simulations in C: *N* = 10000, *p* = 2, *γ* = 0.2%.

We proceed by studying how the maximal fraction of shared neurons varies as a function of the memory load *α* = *P/N* (Fig. 2B). As the load increases, we observe that the maximal fraction of shared neurons decreases, but the change is modest. This weak dependence on the load is robust against two variations of the network where (i) self-interaction of neurons is excluded; or (ii) the *P*−2 background patterns are also overlapping in pairs, e.g., pattern 3 is overlapping with pattern 4, 5 with 6, etc. For both modifications, the mean-field equations look slightly different (Methods) but neither modification leads to a significant change of the maximal fraction of shared neurons *c*_max_ (Fig.2A). In a network that has stored a total of *P* memory engrams, the maximal fraction of shared neurons could potentially depend on the group size *p* of patterns that are all overlapping with each other. So far we have considered *p* = 2. We extended the mean-field approach to the case of three and four overlapping patterns (SI) by rewriting and adapting Eq. (10). Again we find that the maximal overlap is not significantly influenced by the group size *p* of overlapping patterns (Fig. 2B). The group size *p* can be large provided that the total number of patterns *P* does not exceed the memory capacity of the network.

In summary, we found a maximal fraction *c*_max_ of shared neurons beyond which the retrieval of single concepts is no longer possible. The value of *c*_max_ depends on frequency-current curve of neurons.

### What is the minimal fraction of shared concept cells to encode associations?

We find that a symmetric double-retrieval state exists where two concepts are recalled at the same time (Fig 1D, top), even if the fraction of shared concept cells is at chance level. This co-activation of two unrelated concepts could be an artifact of the model considered so far.

In order to check whether our findings in Figs. 1 and 2 are generic, we added to the network the effect of inhibitory neurons by implementing a negative feedback proportional to the overall activity of the *N* neurons in the network. Inhibitory feedback of strength *J*_0_ *>* 0 causes competition during recall of memories. We find that for *J*_0_ = 0.5, each of the two concepts can be recalled individually, but simultaneous recall of both concepts is not possible if the fraction of shared concept cells is at chance level (Fig 3A). If we increase the fraction of shared concept cells above *c* = 5%, then individual as well as simultaneous recall of the two associated memories becomes possible (Fig 3B). The effect becomes even more pronounced at *c* = 20% (Fig 3C). If the fraction of shared neurons reaches a high value of *c*_max_ = 50%, then the separate retrieval of the two individual concepts is no longer possible, indicating that the two concepts have merged into a single one (Fig. 3D). Thus, in the presence of inhibition of strength *J*_0_, we find that the fraction *c* of shared neurons must be in a range *c*_min_(*J*_0_) *< c < c*_max_(*J*_0_) to enable individual as well as joint recall of associated concepts. For *J*_0_ *<* 0.5, the minimal fraction *c*_min_(*J*_0_) is at chance level and for *J*_0_ = 0.5 at *c*_min_ = 5%.

**Figure 3.**
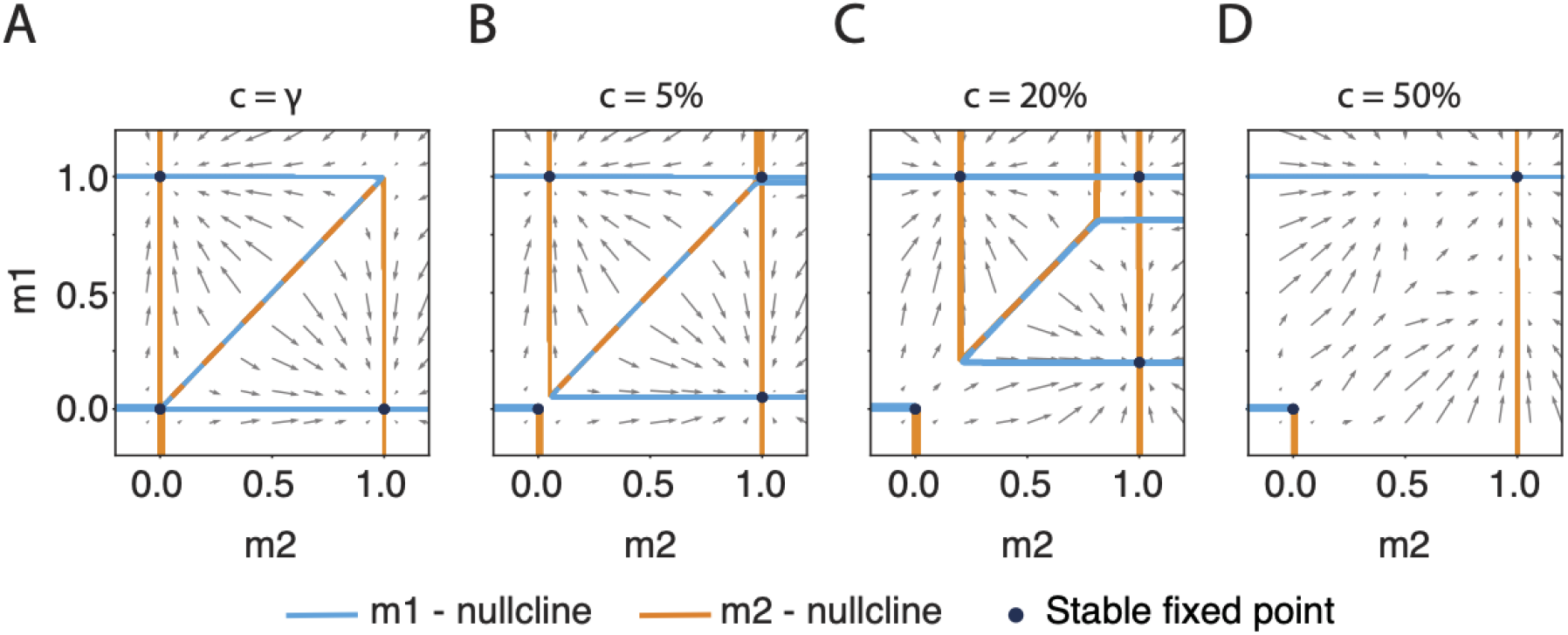
The existence of a symmetric double-retrieval state requires a fraction of shared neurons above chance level in the presence of global inhibition. Four phase-planes showing the stable fixed points in presence of global inhibition, for a fractions of shared neurons **A** *c* = *γ*, **B** *c* = 5%, **C** *c* = 20%, **D** *c* = 50%. On the diagonal, nullclines lie nearly on top of each other (dashed line). Parameters: 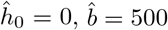, *r*_max_ = 1, *α* = 0, *γ* = 0.2%, *J*_0_ = 0.5.

### Association chains

Neurons shared between memory engrams have been proposed to be the basis for the recall of a memorized list of words [9–12, 26]. In order to translate this idea to chains of associated concepts (Fig. 4A), we follow earlier work [9–12, 26] and add two ingredients to the model of the previous subsection. First, the strength of global inhibitory feedback is now periodically modulated by oscillations mimicking Hippocampal oscillatory activity. The oscillations provide a clock signal that triggers transitions between overlapping concepts. Second, we add to each neuron *i* an adaptation current *θ*_*i*_(*t*) in order to prevent the network state to immediately return to the previous concept. With this extended model, the network state hops from one concept to the next (Fig. 4B). Transitions are repeated, but after some time the network state returns to one of the already retrieved memories, leading to a periodic cycle of patterns [9] (Fig. 4B). In network simulations where concepts are represented by sparse memory engrams (*γ* = 0.2%), we allow a subgroup of *p* = 2, 4 or 16 memory engrams to share a fraction of neurons of *c* = 20%. Because the number of shared concept cells is identical between all pairs of concepts within the same subgroup, the order of the recalled concepts depends on the initial condition. If the subgroup of overlapping engrams is small (*p* = 2, 4), all memory items are retrieved, while for a large group of overlapping engrams (*p* = 16) the cycle closes once a subgroup of the overlapping memory engrams has been retrieved (Fig. 4B).

**Figure 4.**
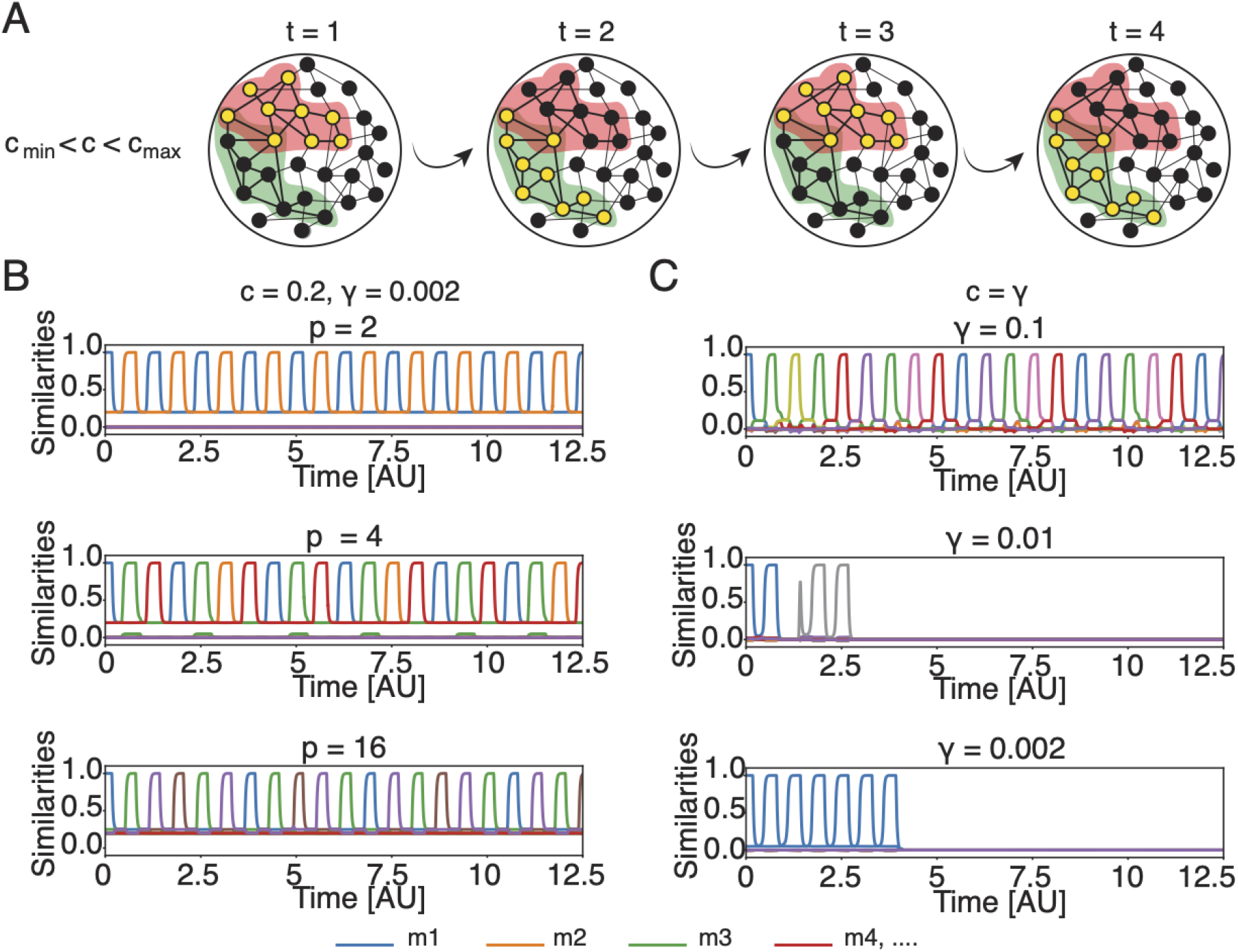
Chain of associations requires shared concepts cells. A) Schematic of a chain of association cycling between two concepts. Assignment of cells to assemblies is indicated by halos’ color. Filled black circles indicate inactive neurons and filled yellow circles indicate active neurons. The schematics corresponds to the top plot of panel B. B) Full network simulation for engrams overlapping above chance level (*c* = 20% *> γ*) with low sparsity (*γ* = 0.2%). Each line corresponds to the similarity *m*^*µ*^ with one of the stored memory engrams as a function of time. A subgroup of *p* engrams is overlapping (top to bottom: *p* =2,4,16. If the network state is initialized to retrieve one of the overlapping concepts, other concepts within the subgroup are retrieved later. C) Same as in B, but memory engrams are independent (*c* = *γ*) and only share cells by chance. By decreasing their mean activity *γ*, the retrieval dynamics of a chain of memories is disrupted. The match between mean-field theory and simulations is shown in Fig. S5. Parameters: *N* = 10 000, *P* = 16, 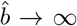, *τ*_*θ*_ = 1.125 s, *T* = 3.75 ms, 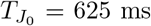, *τ* = 25 ms, *r*_max_ = 40 Hz.

In previous studies [9–12, 26], each memory engram involved a large fraction (*γ* = 10%) of neurons so that transitions could rely on the number of units shared by chance. However, given that the value of the sparsity in MTL is much smaller (*γ* ∼0.23%), it is natural to ask whether the number of neurons shared by chance (*c* = *γ*) is sufficient to induce a sequence of memory retrievals. Our simulations indicate that this is not the case (Fig. 4C). Thus, in a network storing assemblies with a realistic level of sparsity *γ* ∼0.2%, memory engrams with a fraction of shared neurons above chance level are necessary for the retrieval of chains of concepts.

To better understand the role of overlaps between engrams for the formation of association chains, we extend the mean-field dynamics to include the global feedback with periodic modulation *J*_0_(*t*). Since simulations indicate that overlaps are necessary, we want to estimate the minimal and maximal fractions of shared neurons required to enable association chains. Because, in our model, the periodic modulation of the global inhibition strength *J*_0_(*t*) is slow, we consider the mean-field dynamics and the corresponding phase portraits quasi-statically at the two extreme cases, where *J*_0_ is at its maximum and where *J*_0_ is at its minimum. For our parameter setting, when *J*_0_(*t*) is clamped at its *minimum*, the network possesses three stable states: the resting state and the two single retrieval states (Fig. 5B, left). For a successful association chain, we need that concepts can be retrieved separately. The fraction of shared neurons, 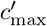, that makes the two single retrieval states disappear therefore sets the upper bound of the useful range of *c*. The parameter 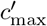 is analogous to *c*_max_ in the previous section, but evaluated in the presence of perdiodic inhibition.

**Figure 5.**
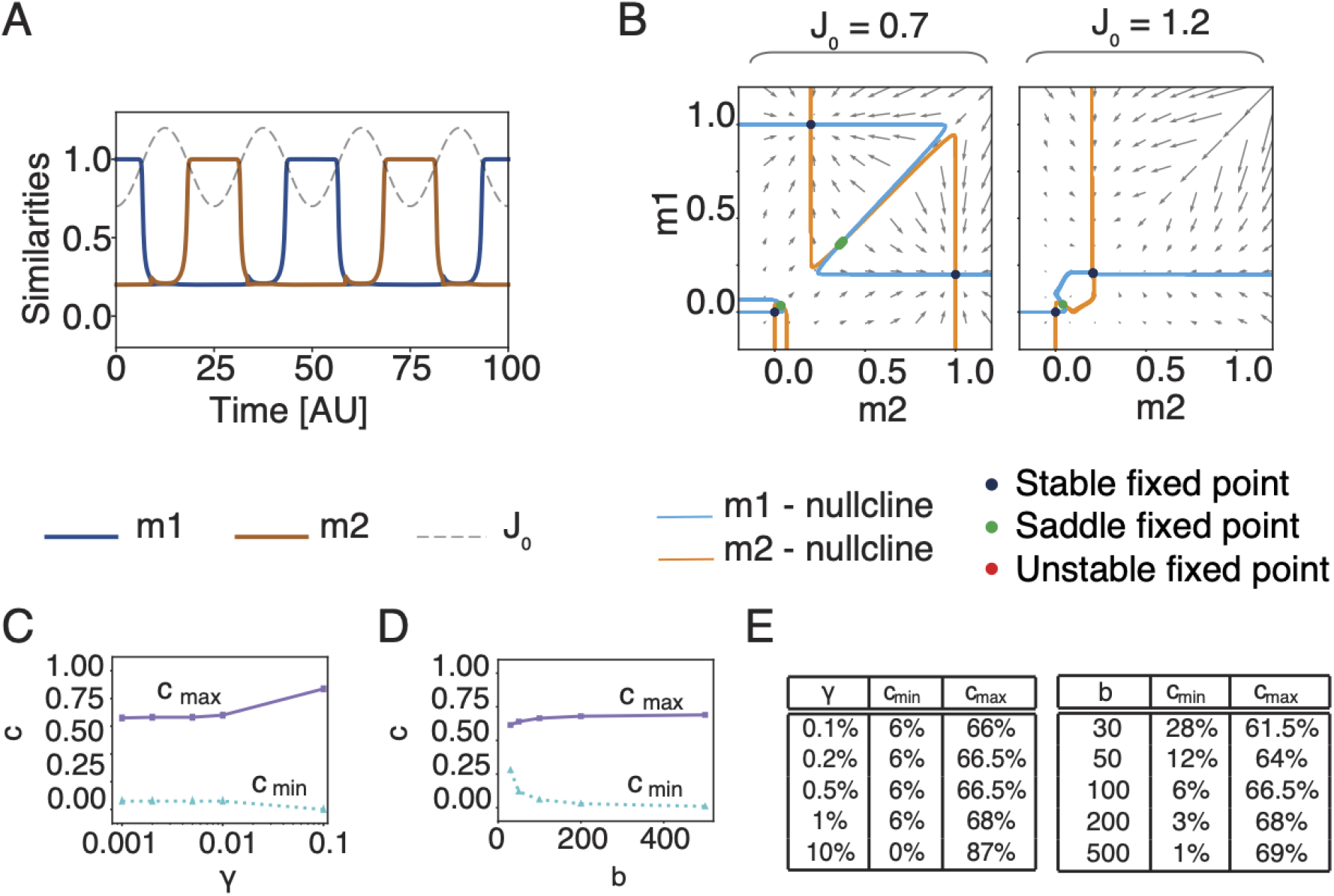
Dependence of association chains on sparsity and neuronal parameters. A) Dynamical mean-field solutions for *m*^1^ and *m*^2^ in the case of two correlated patterns. The grey dashed line shows the modulation of *J*_0_(*t*). B) Phase planes corresponding to the minimum (*J*_0_ = 0.7) and maximum (*J*_0_ = 1.2) value of inhibition in the case of two associated patterns. C) Minimal and maximal fraction of shared concept cells as a function of the sparsity *γ* and D) of the steepness *b*. E) Table with the values of C and D. Parameters (unless specified): *γ* = 0.2%, 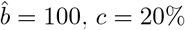, *τ*_*θ*_ = 1.125 s, *T* = 3.75 ms, 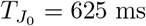, *θ*_*i*_ = 0 for every *i*.

Next, we consider the situation when the global inhibition is clamped at its *maximum* and find the minimal fraction such that the system has, besides the resting state, a second fixed point for *m*^1^ = *m*^2^ *>* 0 where the assemblies of both the previous and next concept are simultaneously active at low firing rates. Since this state is necessary to enable the transition, we call it the *transition state*. If the transition state is present, the network could, once global inhibition decreases, either return from the transition state to the previous concept, or jump to the next one (Fig. 5B, right side). However, in the presence of adaptation (which is not included in the phase plane picture of Fig. 5), the transition to the next concept is systematically favored because neurons participating in the assembly of the earlier concept are fatigued. The existence of the transition state is a necessary condition for the formation of temporal association chains. Thus, the lower bound of the fraction of shared neurons 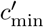 is the smallest overlap such that the transition state exists. Since in the mean-field limit, the transition state appears only for *c > γ*, a fraction of shared neurons above chance level is needed to allow the hopping between concepts. In Fig. 5C-E we show the dependence of the maximal and minimal fraction of shared concept cells upon the sparsity *γ* and the steepness *b*: in both cases the dependence is not strong, but sparser networks lead to a slightly smaller range of the admissible fraction *c* of shared neurons supporting association chains. Importantly, the minimal fraction of shared neurons necessary for association chains is significantly above the fraction of neurons that are shared by chance. We find that for a suitable choice of neuronal and network parameters, association chains are possible for realistic values of *γ* and *c* as measured in human MTL. This suggests that, in principle, associations could be implemented as sequences of transitions if the number of shared neurons is above *c*_min_.

In conclusion, we have shown the need for overlaps between memory engrams – equivalent to a number of shared concept cells significantly above chance level – to explain free memory recall as a chain of associations in recurrent networks such as the human CA3 where each engram involves only a small fraction of neurons.

### How does a network embed groups of overlapping memories?

In our discussion on shared concept cells, we have so far mainly focused on neurons that are shared between a single *pair* of memory engrams such as one place and one person. However, humans are able to memorize many different persons and places, some memories forming subgroups of associated items, others not. In order to compare our network model with human data we therefore need to encode *several* subgroups of *two or more* of overlapping memory engrams in the same network of *N* neurons. Based on the results of the previous sections, we wondered whether we can explain the experimental distribution of the number of concepts a single neuron responds to. We find that imposing the fraction *c* of shared concept cells between *pairs* of concepts, does not predict uniquely how many neurons are used if a given number of memory engrams is embedded in a network. Therefore, imposing *c* as a target number of shared concept cells while encoding multiple concepts is not sufficient to predict whether a given neuron responds to 3 or 5 different concepts. The question then is: to how many concepts does a single neuron respond if several groups of overlapping engrams have been embedded in the network?.

To study this question we consider three different algorithms that all construct memory engrams of 200 neurons per memory with a pairwise overlap of 8 neurons in a network of 100,000 neurons, i.e., *γ* = 0.2% and *c* = 4% (Fig. 6B). When we embed subgroups of 16 engrams with identical numbers of pairwise shared neurons, then an iterative overlap-generating model needs about 2400 neurons out of the 100,000 available neurons, whereas two different hierarchically organized algorithms need about 2400 or 3000 neurons, respectively. In order to understand which of the three algorithms explains experimental data best, we quantify the predictions of the three algorithms under the assumption that not just one, but several subgroups of patterns are embedded in the same network and compare the predictions with experimental data using a previously published dataset of human concept cells [6].

**Figure 6.**
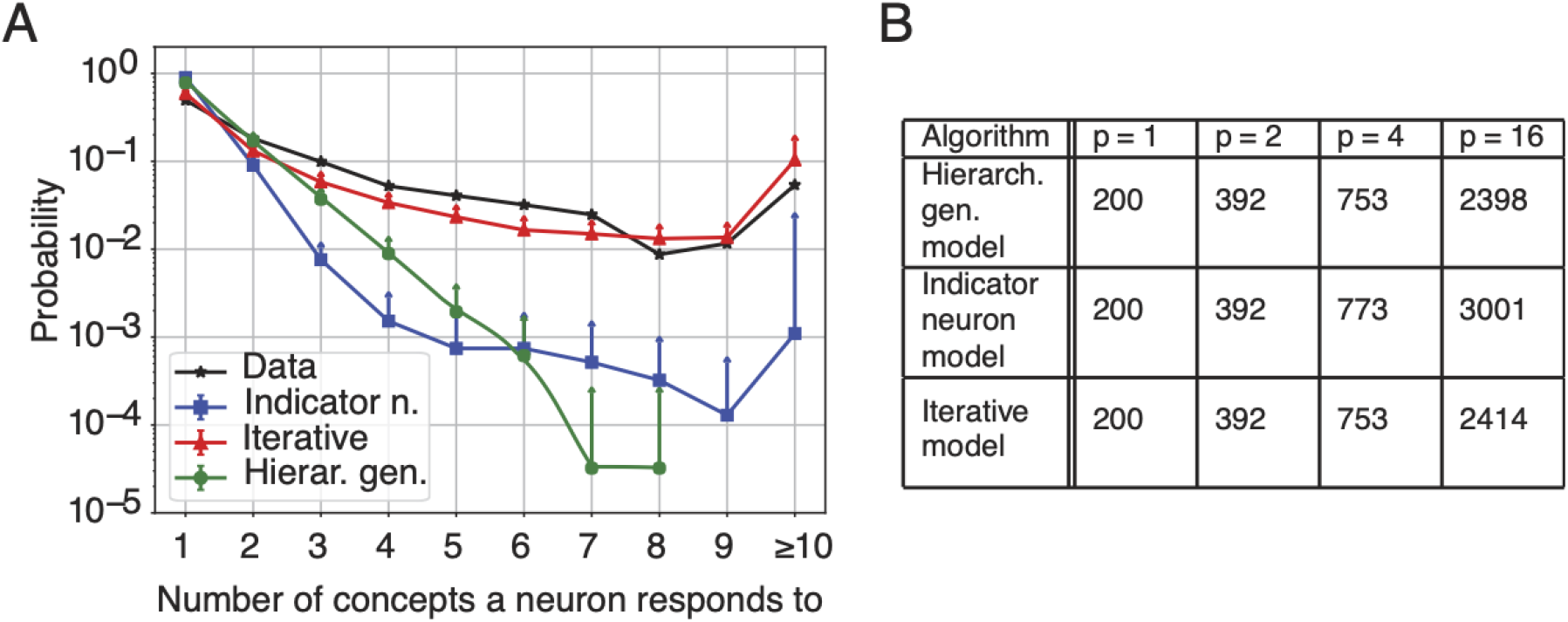
A single neuron responds to several concepts. A) Probability that a neuron responds to a given number of concepts: comparison between data and 3 different algorithms: the hierarchical generative model and the indicator neuron model, which both build overlapping engrams in a hierarchical way, and the iterative overlap-generating model which is a non-hierarchical algorithm. Each algorithm was run 40 times to generate the mean and error bars (only upward bars are displayed, corresponding to one standard deviation). B) For each of the three algorithms, we generated three subgroups of patterns containing *p* = 16, *p* = 4, or *p* = 2 patterns, respectivley, as well as an isolated pattern (*p* = 1). The table gives the expected total number of active neurons in each subgroup in a neural network of 100 000 neurons if patterns have sparsity *γ* = 0.2% and a pairwise fraction of shared neurons *c* = 4%.

The dataset contains the activity of 4066 neurons recorded from the human MTL during the presentation of several visual stimuli. We can extract the experimental probability that a single neuron responds to exactly *k* different concepts (Fig. 6A, black stars). From the probability distribution, we observe the existence of neurons responding to a large number of concepts (10 or more), but also a sizable fraction of neurons that respond to 5 or 6 different concepts. We will refer to those neurons as multi-responsive neurons.

To describe the data, we take into account the size and number of subgroups used in the experimental stimulation paradigm (SI). We find that only the iterative overlap-generating model fits the data (Fig. 5.C), i.e., it is the only one that predicts the correct probability of multi-responsive neurons. Since the iterative overlap-generating model is not based on a hierarchical generation of patterns, this suggests that the MTL encodes large subgroups of memory engrams in a non-hierarchical way, in agreement with earlier papers [6].

### Robustness to heterogeneity

Because biological neural networks present different forms of heterogeneity, we have checked our model’s robustness to (i) the heterogeneity of frequency-current curves and (ii) dilution of the number of synaptic connections.

In the experimental data set, each neuron is characterized by different baseline firing rates and maximal rates in response to the preferred stimulus. We therefore introduce in our model heterogeneous frequency-current curves characterised by a minimum and a maximum firing rates (*r*_min_)_*i*_ and (*r*_max_)_*i*_ respectively and renormalize the network dynamics appropriately (Methods). Despite the heterogeneity, simulations indicate that memory recall with heterogeneity is nearly indistinguishable from that without (compare Figs 7A and 2C and 7B and 4B).

**Figure 7.**
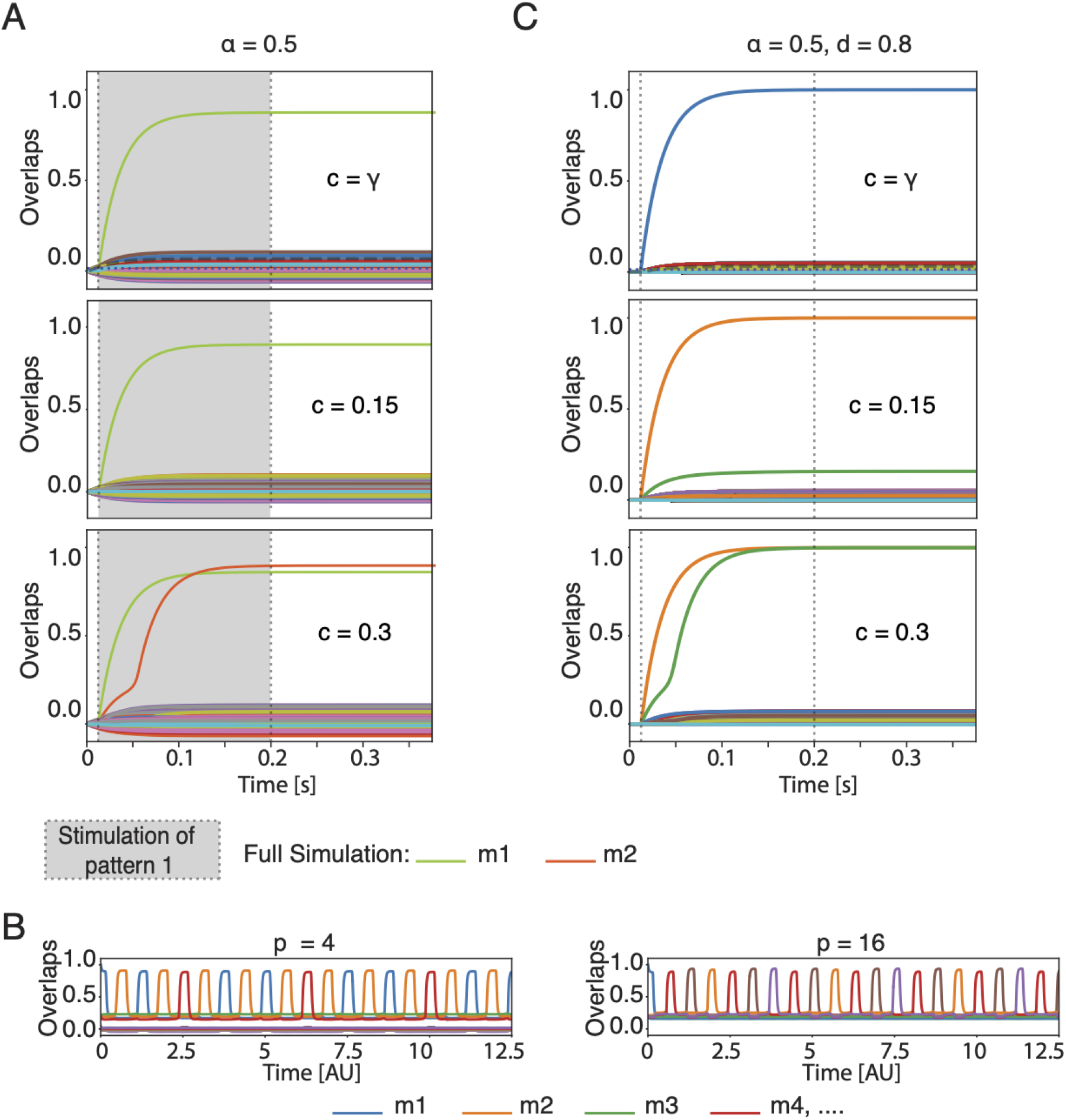
The model is robust to heterogeneity of frequency-current curves. Full network simulations A) in absence of adaptation, equivalent to Fig. 2C, and B) in presence of adaptation and periodic inhibition. C) The model is robust to the dilutions of the synaptic connections. Full network simulations equivalent to Fig. 2C

Secondly, we allow the weight matrix to be diluted. Whereas so far we have assumed an “all-to-all” connectivity, we now introduce the dilution coefficient *d*, which indicates the fraction of actual synaptic connections compared to the *N* ^2^ potential ones. Importantly, for sparsely connected networks, the theory still contains the parameter *α* for memory load, except that *α* is redefined to *α* = *P/M* where *M* is the mean number of connections per neuron (see SI for details). Simulations in Fig. 7C show that the model is robust for *d* = 0.8, i.e. after dropping 20% of all possible synaptic connections and an appropriate rescaling of the average connection strength.

## Discussion

Our results bridge observations and theories from four different fields: first, experimental observations in the human MTL [1, 3, 6, 27, 28]; second, experimental observations of memory engrams [18, 19]; third, the theory of association chains used to explain free memory recall [9–12]; and fourth the classic theory of attractor neural networks [13, 25]. Our main result is that, in networks were concepts are encoded by sparse assemblies, the number of shared concept cells must be above chance level but below a maximal number in order to enable a reliable encoding of associations. With 4-5% overlap between memory assemblies as reported in the human MTL [6], association chains are possible for a range of parameters of frequency-current curves. Our work extends the classical mean-field formalism [15] to memory engrams that exhibit pairwise overlap, both in a static and chain-like retrieval setting.

While sparsity limits the number of concept cells shared by chance, Hebbian learning could induce sharing of concept cells between a small number of specific memories engrams [6]. The existence of a maximal fraction of shared neurons implies that Hebbian learning must work with an intrinsic control mechanism so as to avoid unwanted merging of separate concepts.

Association chains could form the basis of a “stream of thought” where the direction of transitions from one concept to the next is based on learned associations. Our oscillatory network dynamics is inspired by the model of Romani, Tsodyks and collaborators [9–12]. Even though in the Romani- Tsodyks model memory engrams are independent, finite size effects make some pairs of engrams share neurons above chance level which enables sequential recall in the presence of a periodic background input. We find that in large networks with sparse coding level (*γ* ≈0.23%), neurons shared by chance are not enough to reliably induce the retrieval of a chain of concepts. Sequential memory retrieval is possible only for overlaps larger than chance, potentially representing associations learned during real-life episodes. Instead of transitions triggered by oscillations, transitions could also triggered by two adaptation mechanisms that act on different time scales without the need of periodic inhibition [29–31].

Attractor networks with sparse patterns [17] and random connectivity [32] are suitable candidate models for biological memory because they present two features: (i) memory retrieval after stimulation with a partial cue and (ii) sustained activity after a stimulus has been removed. One of the points of critique of attractor networks, traditionally analyzed with the replica [33] or cavity [34, 35] method, has been the unrealistic assumption of symmetric connections. However, the derivation used here, based on dynamical systems arguments [36], can easily be generalized to the case of asymmetric connectivity.

The maximal number of patterns that can be stored in an attractor neural networks has attracted a lot of research [15, 17, 37]. However, does the hippocampus actually operate in the regime of high memory load? Even though we do not believe that hippocampus stores words, we may estimate a rough upper bound for the load *α* = *P/M* in the area CA3 of the hippocampus from the number of words a native English speaker knows (which is about *P* =30’000 according to The Economist, Lexical Facts) and the number of input connections per neuron (which is about *M* =30’000 [38]). Hence we estimate an upper bound of *α* about 1 if concepts are stored in area CA3 – and our theory captures such a high load.

The maximal fraction of neurons which two concepts can share before they effectively merge into a single concept mainly depends on two dimensionless parameters: the rescaled threshold *ĥ* = *h*_0_*/*(*Ar*_max_) and the rescaled steepness 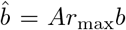. Since these parameters have so far not been estimated for the human CA3 area of the hippocampus or for the MTL in general, we checked parameters of the frequency-current curve of Macaque inferotemporal cortex [24], for which we find *c*_m*ax*_ = 34%.

Finally, by comparing the experimental measured number of concepts a neurons responds to and model predictions we find that the iterative overlap-generating model can predict the number of multi-responsive neurons quite accurately. The algorithm of how to build overlapping engrams plays a key role in fitting the experimental data and confirms the idea that memory engrams in the hippocampus are not hierarchically organised.

## Methods

We consider an attractor neural network of *N* rate units with firing rates *r*_*i*_, in which *P* memory engrams are stored. Each engram *µ*, 1 ≤ *µ* ≤ *P*, is given by a binary random pattern 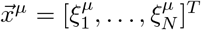, wher 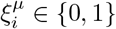 are Bernoulli random variables with mean 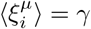. Here and in following,⟨ · ⟩ indicates expectation over the random numbers 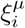 that make up the patterns. Each neuron follows the rate dynamics of Eq. (1), where the synaptic weight from neurons *j* to neurons *i* is defined as [17, 24]

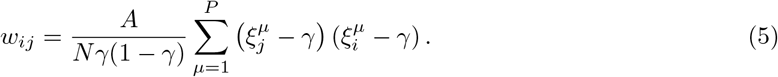

Here, the constant *A* can be interpret as the global scale of “connection strength”. For independent patterns, the synaptic weight *w*_*ij*_ has mean zero, (*w*_*ij*_) = 0, and variance 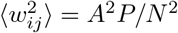.

### Model without adaptation and global feedback

For deriving the results in Figs. 1 - 2, the total input driving neuron *i* is

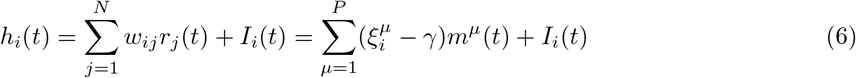

where *I*_*i*_ is the external input. The similarity measure (also called “overlap” in the attractor network literature) *m*^*µ*^ measures the similarity (correlation) of the current network state with pattern *µ*; cf. Eq. (2). In Figs. 1C,2C and 6 the external input 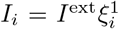 is positive during stimulation for all neurons that belong to the assembly of pattern *µ* = 1.

### Model with adaptation and global inhibitory feedback

For Fig. 4 of Results, we added adaptation and a global inhibitory feedback to the model as described in previous studies [9–12]. Specifically, we add two negative feedback terms to the input potential:

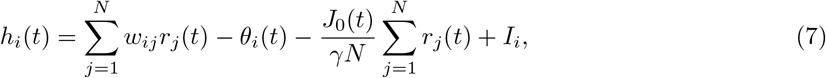

First, the variable *θ*_*i*_(*t*) models neuron-specific firing-rate adaptation via the first-order kinetics

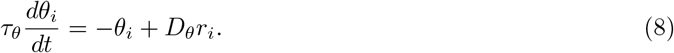

Here, *τ*_*θ*_ is the adaptation time constant and *D*_*θ*_ determines the strength of adaptation. Note that this adaptation model with a hyperpolarizing feedback current is equivalent to a model in which adaptation is implemented as an increase in the threshold *h*_0_ + *θ*_*i*_(*t*).

Second, the global inhibitory feedback term proportional to *J*_0_(*t*) (third term in (7)) provides a clock signal that triggers transitions betweeen attractors. Importantly, inhibition proportional to the summed activity of the network units penalizes network configurations with many active neurons and therefore reduces stability of the double-retrieval state where two memories are recalled together. Here, the strength *J*_0_(*t*) of the global feedback is modulated periodically between values 0.7 and 1.2 with a sinusoidal time course of period *T*_*J*_0 that sets the time scale of transitions between memories. Note that the model without adaptation and global feedback is a special case of the full model by setting *D*_*θ*_ = 0 and *J*_0_(*t*) ≡ 0. For the results of Fig. 3, *J*_0_ is a constant parameter and *D*_*θ*_ = 0.

### Mean-field equations for two overlapping patterns

Overlap between two engrams is implemented as two patterns with a non-zero Pearson correlation coefficient. Without loss of generality, we take patterns 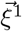 and 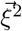 to be correlated, while all other *P* – 2 patterns are independent. We define the correlation *C* between the two patterns as the Pearson correlation coefficient (covariance/variance):

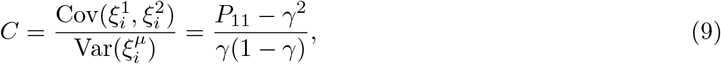

where 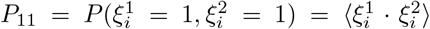 is the joint probabili y f a n eu ron to be selective to both patterns. We generate correlated patterns with mean activity 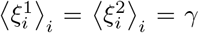 and correlation coefficient *C*, using the procedure described in SI. The fraction *c* of shared neurons is related to *C* by the identity *c* = *C*(1−*γ*) + *γ*.

We are interested in the retrieval dynamics of the correlated patterns 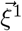 and 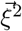. In the limit of large network size, *N* → ∞, this dynamics is given by the mean-field equations for *m*^1^ and *m*^2^ in Eq. 3. For the general case, where the load *α* = *P/N* does not necessarily vanish in the limit *N* → ∞, the functions *F*_*µ*_, *µ* = 1, 2, are given by

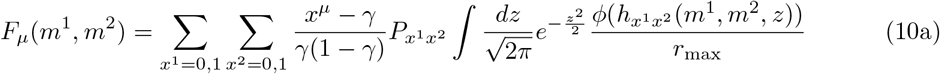

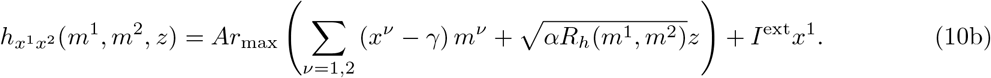

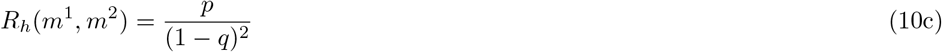

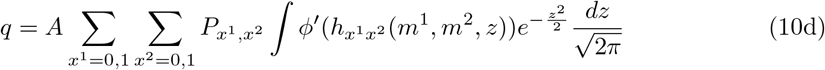

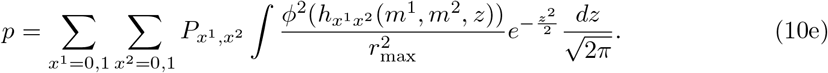

Note that for vanishing load, *α* = *P/N* → 0 in the limit *N* → ∞, we can set *α* = 0 in Eq. (10b) and 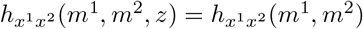 in Eq. (10a), which drastically simplifies the expression for *F*_*µ*_ in closed analytical form (no integral). For details see SI.

A completely analogous procedure can be used to generate more than two correlated binary patterns. In Fig. 2B we generate a subgroup of *p* = 2, 3, 4 correlated patterns and compute their maximal correlation by solving the system of equations that is analogous to (10). To generate Fig. 4C-D we use the mean-field dynamics in the presence of adaptation and global feedback, analogously to Eq. (10). All details are provided in the SI.

### Excluding self-interaction

In order to make the network more biolocally plausible, we can consider that a neuron does not send direct input to itself. The effect of excluding the self-interaction term in Eq. (6) is captured by a correction term to be included on the right-hand-side of Eq. (10b) [36]:

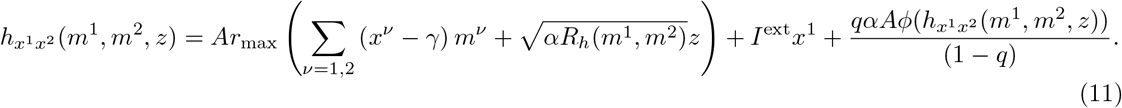

### Overlapping background patterns

In Fig. 2B we explore the possibility that the maximal fraction of shared neurons *c*_max_ might be influenced by the presence of Pearson’s correlation between pairs of background patterns. Moreover, the assumption that there are many subgroups of overlapping memory engrams seems more biologically plausible. If we let the background patterns to be overlapping in subgroups of 2 patterns each, the variable *R* in the mean-field equations of Eq. (10) needs to be replaced by

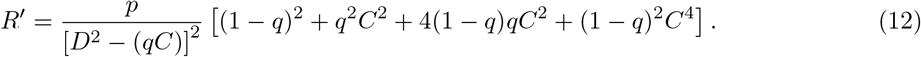

The detailed derivation is provided in the SI.

Furthermore, Eq. (12) can be extended to the case in which background patterns share correlation *C* between non-overlapping subgroups of exactly *p* patterns (the complete derivation is provided in the SI).

### Algorithms to generate overlapping patterns

In this section we describe how a single subgroup of *K* overlapping patterns with sparseness *γ* and overlap *c*is created according to three different algorithms. Details and the theoretical probability distribution associated to the algorithms are given in the SI.

### Hierarchical generative model

We start by creating a “parent” pattern which is not part of the subgroup. The parent pattern has sparseness *λ* = *γ/c*, i.e., 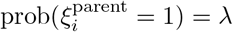. We proceed to create the actual patterns by copying the ones of the parent pattern with probability *c*, while the zeros stay untouched: 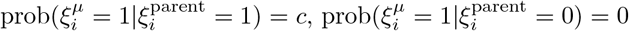.

### Indicator neuron model

We generate with probability *λ* a small subset of indicator neurons. This subset gives a parent pattern of indicator neurons: 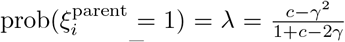. In a network of *N* neurons, we select *n*_*ind*_ = *λN* neurons as indicator neurons. To create a pattern *µ*, we flip the bits of the parent pattern with probability *ϵ* (i.e. a bit is flipped either from 0 to 1 or from 1 to 0 with probability *ϵ*.) The desired values of *λ* and *ϵ* are obtained by expressing the correlation coefficient *C* and the sparseness *γ* as a function of them and reversing the formula.

### Iterative model

We start by generating the first pattern with *γN* active neurons. Neurons that have not been selected for constructing any pattern are classified as “untouched neurons”. For the construction of each of the following patterns, from 2 to *p* (where *p* is the number of correlated patterns), we select randomly *cN* neurons from each of the already created patterns. While building pattern from 2 to *p*, we count the amount of already shared neurons between the pattern under construction and the one we are picking the shared units from. Therefore and take this into account, by picking *cN* minus the number of already shared neurons. Finally we pick the remaining neurons to reach the target of *γN* active neurons from the untouched ones.

### Experimental data

The experimental dataset of Fig. 6 comes from a previous publication [6]. The data was recorded from patients implanted with chronic depth electrodes in the MTL for the monitoring of epileptic seizures. Micro-wires recorded the localized neural activity; spike detection and sorting allowed to identify the activity of 4066 single neurons. During recordings, patients were shown different pictures of known people and places repeated several times. For each neuron, the stimuli eliciting a response were identified using a statistical criterion based on the modulations of firing rate during stimulus presentation compared to baseline epochs. For additional details on the dataset and data processing we refer to the original publication.

### Heterogeneous frequency-current curves

The frequency-current function of model neurons is neuron-specific and re-written as

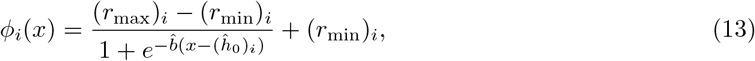

where the values of (*r*_min_)_*i*_ and (*r*_max_)_*i*_ are randomly sampled for each neuron from a Gaussian distribution with mean and standard deviation *µ*_min_, *σ*_min_ and *µ*_max_, *σ*_max_ respectively. The parameter (*ĥ* _0_)_*i*_ is then defined as *h*_0_((*r*_max_)_*i*_− (*r*_min_)_*i*_), where *h*_0_ is a global constant. Finally in the firing rate equation, we re-scale the firing rates as follows:

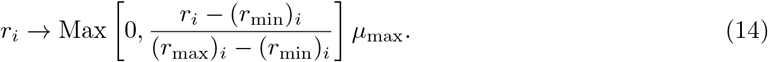

In Fig. 7 we choose the parameters *µ*_min_ = 0 Hz, *µ*_max_ = 1 = 40 Hz, and *σ*_min_ = *σ*_max_ = 4 Hz.

## Acknowledgements

We thanks to Valentin Marc Schmutz, Martin Barry, Alireza Modirshanechi and Johanni Brea for useful comments and discussions. This research was supported by the Swiss National Science Foundation, grant agreement 200020 165538 and by the European Union Horizon 2020 Framework Program under grant agreement no. 785907 (HumanBrain Project, SGA2).

## References

[1] R Quian Quiroga, Leila Reddy, Gabriel Kreiman, Christof Koch, and Itzhak Fried. Invariant visual representation by single neurons in the human brain. Nature, 435(7045):1102, 2005.

[2] Matias J Ison and Rodrigo Quian Quiroga. Selectivity and invariance for visual object perception. Front Biosci, 13:4889–4903, 2008.

[3] Rodrigo Quian Quiroga. Neural representations across species. Science, 363(6434):1388–1389, 2019.

[4] Stephen Waydo, Alexander Kraskov, Rodrigo Quian Quiroga, Itzhak Fried, and Christof Koch. Sparse representation in the human medial temporal lobe. Journal of Neuroscience, 26(40):10232– 10234, 2006.

[5] Matias J Ison, Rodrigo Quian Quiroga, and Itzhak Fried. Rapid encoding of new memories by individual neurons in the human brain. Neuron, 87(1):220–230, 2015.

[6] Emanuela De Falco, Matias J Ison, Itzhak Fried, and Rodrigo Quian Quiroga. Long-term coding of personal and universal associations underlying the memory web in the human brain. Nature communications, 7:13408, 2016.

[7] Hernan G Rey, Emanuela De Falco, Matias J Ison, Antonio Valentin, Gonzalo Alarcon, Richard Selway, Mark P Richardson, and Rodrigo Quian Quiroga. Encoding of long-term associations through neural unitization in the human medial temporal lobe. Nature communications, 9(1):1–13, 2018.

[8] Hernan G. Rey, Belen Gori, Fernando J. Chaure, Santiago Collavini, Alejandro O. Blenkmann, Pablo Seoane, Eduardo Seoane, Silvia Kochen, and Rodrigo Quian Quiroga. Single neuron coding of identity in the human hippocampal formation. Current Biology, 30(6):1152 – 1159.e3, 2020.

[9] Sandro Romani, Itai Pinkoviezky, Alon Rubin, and Misha Tsodyks. Scaling laws of associative memory retrieval. Neural computation, 25(10):2523–2544, 2013.

[10] Stefano Recanatesi, Mikhail Katkov, Sandro Romani, and Misha Tsodyks. Neural network model of memory retrieval. Frontiers in computational neuroscience, 9:149, 2015.

[11] Stefano Recanatesi, Mikhail Katkov, and Misha Tsodyks. Memory states and transitions between them in attractor neural networks. Neural computation, 29(10):2684–2711, 2017.

[12] Michelangelo Naim, Mikhail Katkov, Sandro Romani, and Misha Tsodyks. Fundamental law of memory recall. Physical Review Letters, 124(1):018101, 2020.

[13] John J Hopfield. Neural networks and physical systems with emergent collective computational abilities. Proceedings of the national academy of sciences, 79(8):2554–2558, 1982.

[14] Gérard Weisbuch and Françoise Fogelman-Soulié. Scaling laws for the attractors of hopfield networks. Journal de Physique Lettres, 46(14):623–630, 1985.

[15] Daniel J Amit and Daniel J Amit. Modeling brain function: The world of attractor neural networks. Cambridge university press, 1992.

[16] I Kanter and Haim Sompolinsky. Associative recall of memory without errors. Physical Review A, 35(1):380, 1987.

[17] Mikhail V Tsodyks and Mikhail V Feigel’man. The enhanced storage capacity in neural networks with low activity level. EPL (Europhysics Letters), 6(2):101, 1988.

[18] Susumu Tonegawa, Michele Pignatelli, Dheeraj S Roy, and Tomás J Ryan. Memory engram storage and retrieval. Current opinion in neurobiology, 35:101–109, 2015.

[19] Sheena A Josselyn and Susumu Tonegawa. Memory engrams: Recalling the past and imagining the future. Science, 367(6473), 2020.

[20] César Rennó-Costa, John E Lisman, and Paul FMJ Verschure. A signature of attractor dynamics in the ca3 region of the hippocampus. PLoS Comput Biol, 10(5):e1003641, 2014.

[21] Tom J Wills, Colin Lever, Francesca Cacucci, Neil Burgess, and John O’Keefe. Attractor dynamics in the hippocampal representation of the local environment. Science, 308(5723):873–876, 2005.

[22] Siegfried Bös, R Kühn, and JL van Hemmen. Martingale approach to neural networks with hierarchically structured information. Zeitschrift für Physik B Condensed Matter, 71(2):261–271, 1988.

[23] Vezha Boboeva, Romain Brasselet, and Alessandro Treves. The capacity for correlated semantic memories in the cortex. Entropy, 20(11):824, 2018.

[24] Ulises Pereira and Nicolas Brunel. Attractor dynamics in networks with learning rules inferred from in vivo data. Neuron, 99(1):227 – 238.e4, 2018.

[25] J. J. Hopfield. Neurons with graded response have collective computational properties like those of two-state neurons. Proceedings of the National Academy of Sciences, 81(10):3088–3092, 1984.

[26] Mikhail Katkov, Sandro Romani, and Misha Tsodyks. Effects of long-term representations on free recall of unrelated words. Learning & Memory, 22(2):101–108, 2015.

[27] Rodrigo Quian Quiroga. Concept cells: the building blocks of declarative memory functions. Nature Reviews Neuroscience, 13(8):587–597, 2012.

[28] Rodrigo Quian Quiroga. Plugging in to human memory: advantages, challenges, and insights from human single-neuron recordings. Cell, 179(5):1015–1032, 2019.

[29] Eleonora Russo, Vijay MK Namboodiri, Alessandro Treves, and Emilio Kropff. Free association transitions in models of cortical latching dynamics. New Journal of Physics, 10(1):015008, 2008.

[30] Eleonora Russo and Alessandro Treves. Cortical free-association dynamics: Distinct phases of a latching network. Physical Review E, 85(5):051920, 2012.

[31] Athena Akrami, Eleonora Russo, and Alessandro Treves. Lateral thinking, from the hopfield model to cortical dynamics. Brain research, 1434:4–16, 2012.

[32] Daniel J Amit and Nicolas Brunel. Model of global spontaneous activity and local structured activity during delay periods in the cerebral cortex. Cerebral cortex (New York, NY: 1991), 7(3):237–252, 1997.

[33] Daniel J Amit, Hanoch Gutfreund, and Haim Sompolinsky. Information storage in neural networks with low levels of activity. Physical Review A, 35(5):2293, 1987.

[34] Marc Mézard, Giorgio Parisi, and Miguel Virasoro. Spin glass theory and beyond: An Introduction to the Replica Method and Its Applications, volume 9. World Scientific Publishing Company, 1987.

[35] Maoz Shamir and Haim Sompolinsky. Thouless-anderson-palmer equations for neural networks. Physical Review E, 61(2):1839, 2000.

[36] Masatoshi Shiino and Tomoki Fukai. Self-consistent signal-to-noise analysis and its application to analogue neural networks with asymmetric connections. Journal of Physics A: Mathematical and General, 25(7):L375, 1992.

[37] Daniel J Amit, Hanoch Gutfreund, and Haim Sompolinsky. Storing infinite numbers of patterns in a spin-glass model of neural networks. Physical Review Letters, 55(14):1530, 1985.

[38] Per Andersen, Richard Morris, David Amaral, Tim Bliss, and John O’Keefe. The hippocampus book. Oxford university press, 2006.

