## Supplementary Information for "When shared concept cells support associations: theory of overlapping memory engrams"



#### Summary

Experimental studies measure the fraction of shared memory cells $c$. Such fraction of shared neurons can be interpreted as the probability that a neuron that already responds to one concept also responds to a second one. Formally, the fraction of shared neuron is the *conditional* probability $\text{Prob}(x^2 = 1|x^1 = 1)$, where $x^\mu$ are binary variables that indicate whether a neuron belongs to the cell assembly representing concept mu ($\xi_i^\mu = 1$), or not ($\xi_i^\mu = 0$). In other words, $c$ is the fraction of shared neurons *relative* to the total number of neurons in the assembly under consideration. We define the overlap n between two cell assemblies as the absolute number of neurons that are shared between two cell assemblies $\mu$ and $\nu$: $n = \sum_{i=1}^N \xi_i^\mu \xi_i^\nu = c\gamma N$, where $\gamma$ is the fraction of neurons that take part in a given memory assembly. Note that $c\gamma$ is the *joint* probability that a neuron belongs to two memory assemblies: $P_{11} = \text{Prob}(x^2 = 1, x^1 = 1)$. In the case of two *independent* random patterns $\vec{\xi}^\mu$, $\vec{\xi}^\nu$, the probability that a neuron belongs to both assembly $\mu$ and $\nu$ is $\gamma^2$. We refer to this scenario as the chance level. A chance level, the number of shared neurons (overlap) is $n = \gamma^2 N$ and, hence, the fraction of shared neurons is $c = \gamma$.

From a probabilistic point of view, the fraction of shared neurons c is uniquely determined by the Pearson's correlation coefficient $C$:

$$C = \frac{\text{Cov}(\xi_i^1, \xi_i^2)}{\text{Var}(\xi_i^\mu)} = \frac{P_{11} - \gamma^2}{\gamma(1-\gamma)}, \tag{1}$$

The fraction of shared neurons $c$ can be related to the Person's correlation coefficient $C$ uniquely: $c = n/(\gamma N) = \gamma + (1-\gamma)C$. In the derivation of the derivation of the mean-field equations for overlapping memory engrams, we will refer to C simply as the correlation.


#### Maximal fraction of shared neurons between memory engrams

In the METHODS, we introduce the concept of the critical fraction of shared neurons as the fraction of shared neurons below which each pattern has a separated basin of attraction and above which the basins of attraction merge. The latter means that memories are indistinguishable. As mentioned in the introduction each fraction of shared neurons corresponds to a specific correlation value, so we can equivalently speak of critical correlation between memory patterns. In order to numerically compute the critical correlation, we use the bifurcation diagram in Fig. 1SB: the fixed points in the phase-plane are projected on the $m^1$-axis and their positions are plotted as $C$ increases. From the bifurcation diagram we can extract the value $C_{\max}$ at which the single retrieval states merge with the saddle points and disappear. Thus, at $C = C_{\max}$ we have a saddle-node bifurcation (see the derivation of Eq. (83)).

The value of the critical correlation $C_{\max}$ can be calculated analytically in the limit of infinite steepness $b \to \infty$, vanishing load $\alpha = 0$, vanishing sparseness and load, $\gamma \to 0$ and $\alpha = 0$. This value matches the one extracted from the bifurcation plot in Fig. 1SC.

In the super critical regime $C > C_{\max}$, the activation of patter one triggers, with some delay, the activation of the correlated pattern 2, as shown in Fig. 1D. The activation delay of pattern 2 can be quantified using the dynamical mean-field in Eq. (41). In Fig. 2SA we show the evolution of the system in the phase plane, before, during and after the stimulation respectively. In Fig. 2SB, we can see the activation of $m^2$ due to the super-critical correlation with $m^1$ as predicted by the mean-field theory. We choose to quantify the time delay between the activation of pattern one and that of pattern two, by comparing the $m^1(t)$ and $m^2(t)$ lines in Fig. 2SB when they cross the value $m^1 = m^2 = \hat{h}_0$: time gap between the two crossing times defines the delay. Indeed, the dimensionless parameter $\hat{h}_0$ marks the point where the $m^2(t)$ curve becomes steeper, or, in the phase-planes, it is close to the ghost of the fixed point corresponding to the single retrieval state.

#### Association chains

We estimate the range of correlations such that association chains are possible. That is, correlation should be large enough to trigger the activation of the next pattern, but not so large that the basin of attraction of single patterns merge. The strength of the global inhibition $J_0(t)$ varies slowly between its maximum and its minimum. Note that, when $J_0(t)$ is clamped at its minimum in Fig. 4SB,D,F,H left hand side, the double retrieving state is not present. The lack of the double retrieving state is a consequence of the introduction of the global inhibitory feedback, which leads to competition between the two assemblies and a winner-take-all response. The value of correlation, $C_{\max}$, that makes the two single retrieval states disappear can be read off from the bifurcation diagram in Fig. 3SA. It sets the upper bound of the useful correlation range and strongly depends on the value of mininal global inhibitory feedback, $\min(J_0)$. When we want to estimate the smallest value of correlation, $C_{\min}$ so that hopping between attractors is possible, we consider the situation when the global inhibition is clamped at its maximum and find the minimal correlation such that the system exhibits a *transition state*. The transition state is visible in Fig. 4SD,H right hand side, but it is not present in Fig. 4SF, since $C = 0$. Thus, the lower bound of correlation $C_{\min}$ is estimated by the left end of the stable diagonal branch of fixed points in the bifurcation diagram (Fig. 3SB).

In Fig. 4S we compare the effect of different sparseness and correlations on the when hopping from one attractor to the next. For very sparse patterns, $\gamma = 0.002$, the transition is sharper (Fig. 4SH) and we observe the same dynamics (Fig. 4SG). While chains of associations are not possible for $C = 0$. Indeed, in Fig. 4SB we assume $\vec{\xi}^1$ and $\vec{\xi}^2$ to be independent ($C = 0$) and the state is never able to leave the basin of attraction of pattern 1. The reason is clear from the phase-planes in Fig. 4SA, when $J_0$ is at its maximum, $m^1$ is decreased in value until a stable fixed point, where still $m^2 = 0$: the second pattern is not even partially activated, and when the inhibition decreases, the system state falls back into the first pattern basin of attraction. At the contrary, when enough correlation is added, any activation of $\xi^1$ implies a partial activation of $\xi^2$. When $J_0$ reaches an high value, the system is pushed in a neutral state where both $m^1 = m^2 \sim C$. At the subsequent decrease of $J_0$, the system might fall in either of the two single retrieval states, but adaptation breaks the symmetry and pushes the system towards the pattern that was not activated yet.

In Fig. 5S we compare full network simulation with dynamical mean-field for $p = 2$ and $p = 4$. The mean-field and the full simulation match.

#### Parameters choice

We have discussed that the critical correlation between patterns depends on two dimensionless parameters: the rescaled threshold $\hat{h}_0 = h_0/(Ar_{\max})$ and the rescaled steepness $\hat{b} = Ar_{\max}b$. While these parameters have so far not being estimated for human Hippocampus, the gain function has been fully characterized for pyramidal neurons of the macaque's IT cortex [1]. We therefore computed the critical correlation for this physiological parameter set. First, in [1] the input is processed by the combination of two sigmoids. We have chosen the parameters of our single sigmoidal gain function such that it fits the combination of sigmoids used in [1]. Moreover, whereas in [1] patterns with $N(0,1)$ Gaussian distributed elements have been considered, in our theory we assumed binary patterns. To match roughly match the two settings, we have chosen $\gamma = N(0,1)(h_0)$, where $h_0$ is estimated from the fitted gain function. We obtained $\gamma = 0.0375$. In Fig. 6SA we show the phase-plane for $C = 0$ (or equivalently $c = \gamma$) and in Fig. 6SB we provide the bifurcation diagram from which the critical correlation is extracted (equivalent to that in Fig. 1SB).

### Methods

In this section, we will present the details and the derivation of both the full network simulations and the mean-field model. We partially repeat information present in the main text, with the goal of providing here a complete and self consistent presentation of the methods used.

#### Model without adaptation and global feedback

In the figures relative to the Section "Maximal fraction of shared neurons between memory engrams" of the main paper and the SI, we consider an attractor neural network of $N$ rate units $r_i$, in which $P$ binary memory patterns are stored $\{\xi_i^\mu\}_{i=1,..,N}^{\mu=1,..,P}$. Each neuron follows the Wilson-Cowan dynamics [2]:

$$\tau \frac{dr_i}{dt} = -r_i + \phi(h_i), \tag{2}$$

where $h_i$ is the input term

$$h_i = \sum_{j \neq i}^{N} w_{ij} r_j + I_i(t) \tag{3}$$

which include the sum over the recurrent input from other neurons and a possible external input $I_i(t)$. The input is filtered with gain function $\phi$, which is chosen to be a sigmoid:

$$\phi(h) = \frac{r_{\max}}{1 + e^{-b(h-h_0)}}. \tag{4}$$

The parameters that define the gain function can be interpreted as follows: $r_{\max}$ is the maximal firing rate, $b$ is the steepness of the gain function and $h_0$ is the bias which is commonly interpreted as firing threshold. While $h_0$ is a hard threshold for $b \to \infty$, at finite $b$ the model exhibits a soft threshold allowing firing activity even below $h_0$. The connection matrix, $w_{ij}$, contains the synaptic weights between neurons $i$ and $j$ is defined as [1]

$$w_{ij} = \frac{A}{N\gamma(1-\gamma)} \sum_{\mu}^{P} \left(\xi_j^\mu - \gamma\right)\left(\xi_i^\mu - \gamma\right) \tag{5}$$

where the constant $A$ can be interpret as the overall connection strength. In order for the weights to to have mean zero, $\langle w_{ij} \rangle = 0$, we subtract the mean activity of patterns $\langle \xi_i^\mu \rangle = \gamma$. In order to measure how close is the network state to recalling a pattern, we defined the similarities $m^\mu$ (also referred to overlaps in theoretical neuro-science) which are a measure of the correlation between the network state and the store patterns:

$$m^\mu(t) = \frac{1}{N\gamma(1-\gamma)r_{\max}} \sum_{j=1}^{N} \left(\xi_j^\mu - \gamma\right) r_j(t). \tag{6}$$

#### Model with adaptation and global inhibitory feedback

For Figs. of sections "Association chains", we added adaptation and a global periodic inhibitory to the model, following previous works [3–6]:

$$h_i = \sum_{j \neq i}^{N} w_{ij} r_j(t) - \theta_i(t) - \frac{J_0(t)}{\gamma N} \sum_{j=1}^{N} r_j(t), \tag{7}$$

The second term $-\theta_i(t)$, is a *local* neuron-specific feedback that models firing-rate adaptation as follows:

$$\tau_\theta \frac{d\theta_i}{dt} = -\theta_i + D_\theta r_i, \tag{8}$$

where $\tau_\theta$ is the adaptation time constant and $D_\theta$ determines the strength of adaptation.

The term $-J_0(t)/(\gamma N) \sum_{j=1}^{N} r_j(t)$ is the *global* inhibition which provides a clock signal that triggers the transitions betweeen attractors. The strength of the global feedback, $J_0(t)$, is modulated periodically in time:

$$J_0 = \frac{1}{2}\left(J_{\max} - J_{\min}\right) \sin\left(\frac{2\pi}{T_{J_0}} t - \frac{\pi}{2}\right) + \frac{1}{2}\left(J_{\max} + J_{\min}\right) \tag{9}$$

#### Non-dimensionalization of the model

The calculations below are considerably simplified if the model is made dimension-free. We take into account that $r_{\max}$ has units of 1/time and the parameter $A$ has units of current $\cdot$ time and measure in the following time in units of $r_{max}^{-1}$ and current input in units of $Ar_{max}$.

**Model without adaptation and global feedback.** Using the dimensionless quantities

$$\hat{h}_i = \frac{h_i}{Ar_{\max}}, \quad \hat{h}_0 = \frac{h_0}{Ar_{\max}}, \quad \hat{b} = bAr_{\max}, \quad \hat{r}_i = \frac{r_i}{r_{\max}} \tag{10}$$

$$\hat{w}_{ij} = \frac{w_{ij}}{A}, \quad \hat{\tau} = \tau r_{\max}, \quad \hat{I}_i(t) = \frac{I_i(t)}{Ar_{\max}}, \tag{11}$$

the dimension-free model without adaptation reads

$$\hat{\tau}\frac{d\hat{r}_i}{d\hat{t}} = -\hat{r}_i + \hat{\phi}(\hat{h}_i), \text{ with } \hat{h}_i = \sum_{j=1}^{N} \hat{w}_{ij}\hat{r}_j + \hat{I}_i \tag{12}$$

with a transfer function $\hat{\phi}(\hat{h}) = 1/\{1 + \exp[-\hat{b}(\hat{h} - \hat{h}_0)]\}$.

**Model with adaptation and global feedback.** Introduction of further dimensionless quantities

$$\hat{\tau}_\theta = \tau_\theta r_{\max}, \quad \hat{\theta} = \frac{\theta}{Ar_{max}}, \quad \hat{D}_\theta = \frac{D_\theta}{A}, \quad \hat{J}_0(\hat{t}) = \frac{J_0(\hat{t}/r_{\max})}{A} \tag{13}$$

leads to the non-dimensional model with adaptation

$$\hat{\tau}\frac{d\hat{r}_i}{d\hat{t}} = -\hat{r}_i + \hat{\phi}(\hat{h}_i), \tag{14}$$

$$\hat{\tau}_\theta\frac{d\hat{\theta}_i}{d\hat{t}} = -\hat{\theta}_i + \hat{D}_\theta \hat{r}_i \tag{15}$$

with input

$$\hat{h}_i = \sum_{j=1}^{N} \hat{w}_{ij}\hat{r}_j - \hat{\theta}_i - \frac{\hat{J}_0(t)}{\gamma N}\sum_{j=1}^{N} \hat{r}_j(t) + \hat{I}_i. \tag{16}$$

To lighten the notation, we will omit the hats on top of the dimensionless quantities for the rest of this paper, while we keep the strict distinction between quantities with and without hat in the Results section.

#### Review of attractor theory

Starting from the overlap definition Eq. (6), we can write equations for the overlaps variables. We first focus on the model without adaptation and global feedback. For this case, we follow an approach well-known in literature [7, 8]. The only time-dependent variable in the definition of $m^\mu$ is the network state $r_j(t)$, so that:

$$\hat{\tau}\frac{dm^\mu}{d\hat{t}} = \frac{1}{N\gamma(1-\gamma)}\sum_{j=1}^{N} \left(\xi_j^\mu - \gamma\right)\hat{\tau}\frac{d\hat{r}_j}{d\hat{t}}. \tag{17}$$

By inserting the expression for the single neuron dynamics Eq. (12) and recognizing the overlap definition Eq. (6), we obtain:

$$\hat{\tau}\frac{dm^\mu}{d\hat{t}} = -m^\mu + F_\mu(m^1, \ldots, m^P). \tag{18}$$

with

$$F_\mu(m^1, \ldots, m^P) = \frac{1}{N\gamma(1-\gamma)}\sum_{j=1}^{N} \left(\xi_j^\mu - \gamma\right)\hat{\phi}\left(\hat{h}_j\right). \tag{19}$$

In this equation, the dependence on the overlaps $m^1, ..., m^P$ is contained in the input term $\hat{h}_j$. From Eq. (12) and by using the definition of the weights $w_{ij}$, Eq. (5), we have

$$\hat{h}_i = \sum_{\mu=1}^{P} (\xi_i^\mu - \gamma)\, m^\mu + \hat{I}_i. \tag{20}$$

In what follows, we are interested in finding equilibrium solutions of Eq. (18), for which $m^\mu = F_\mu(m^1, \ldots, m^P)$. Because we are interested in pattern retrieval, we consider, without loss of generality, the retrieval of pattern 1. To this end, we assume that among all $m^\mu$, only $m^1$ is significantly larger than zero. This network state could be the result of a stimulation in the direction of pattern 1: $\hat{I}_i(t) = \hat{I}(t)\xi_i^1$. Under this assumption we can re-write the input term $\hat{h}_i$ isolating the contribution from $m^1$

$$\hat{h}_i = (\xi_i^1 - \gamma)\, m^1 + \sum_{\mu=2}^{P} (\xi_i^\mu - \gamma)\, m^\mu + \hat{I}(t)\xi_i^1. \tag{21}$$

We call the patterns that are not recalled "background patterns"; in our case, these are all patterns for which $\mu \geq 2$. The second term on the r.h.s of Eq. 21 represents the contribution from the background patterns causing some degree of heterogeneity of the input potential for neurons with the same selectivity to pattern 1. For large $P$, this heterogeneity can be captured by replacing the term $\sum_{\mu=2}^{P} (\xi_i^\mu - \gamma)\, m^\mu$ by a Gaussian random variable with mean zero and variance

$$\sigma^2 = \frac{1}{N} \sum_{i=1}^{N} \sum_{\mu=2}^{P} \sum_{\nu=2}^{P} (\xi_i^\mu - \gamma)\, (\xi_i^\nu - \gamma)\, m^\mu m^\nu = \gamma(1-\gamma) \sum_{\nu=2}^{P} (m^\nu)^2 \tag{22}$$

To obtain the result in Eq. (22), we used the assumption that patterns $\xi_i^\mu$ and $\xi_i^\nu$ are uncorrelated, and the fact that only the term for $\mu = \nu$ survives, in fact, $\left\langle (\xi_i^\mu \xi_i^\nu + \gamma^2 - \gamma\xi_i^\nu - \gamma\xi_i^\mu) \right\rangle_i = \delta_{\mu\nu}\gamma(1-\gamma)$. Here and in the following, the brackets $\langle x_i \rangle$ of a variable $x_i$ denotes the population average $\langle x_i \rangle = \frac{1}{N}\sum_{i=1}^{N} x_i$. In the next passages we compute $(m^\mu)^2, \mu \neq 1$, in the large network limit $N \to \infty$. For $\mu = 2, \ldots, P$, we expand Eq. (19) around $m^\mu = 0$ up to first order in $m^\mu$,

$$F_\mu(m^1, \ldots, m^P) \approx \frac{1}{\gamma(1-\gamma)N} \sum_{j=1}^{N} \left[ (\xi_j^\mu - \gamma)\, \hat{\phi}(\hat{h}_j)\Big|_{m^\mu=0} + (\xi_j^\mu - \gamma)^2\, \hat{\phi}'(\hat{h}_j)\Big|_{m^\mu=0} m^\mu \right]. \tag{23}$$

At equilibrium, $m^\mu = F_\mu(m^1, \ldots, m^P)$, and we thus have

$$m^\mu \left( 1 - \sum_{j=1}^{N} \frac{(\xi_j^\mu - \gamma)^2\, \hat{\phi}'(\hat{h}_j)}{\gamma(1-\gamma)N} \right) = \frac{1}{\gamma(1-\gamma)N} \sum_{j=1}^{N} (\xi_j^\mu - \gamma)\, \hat{\phi}(\hat{h}_j), \qquad \mu \geq 2. \tag{24}$$

On the left hand side of the last expression, we can make some simplification, considering that $\xi_j^\mu$ is uncorrelated with $\phi'(h_j)$, in the $N \to \infty$ limit:

$$\lim_{N\to\infty} \frac{1}{N} \sum_{j=1}^{N} \frac{(\xi_j^\mu - \gamma)^2\, \hat{\phi}'(\hat{h}_j)}{\gamma(1-\gamma)} = \lim_{N\to\infty} \frac{1}{N} \sum_{j=1}^{N} \hat{\phi}'(\hat{h}_j) = \left\langle \hat{\phi}'(\hat{h}_i) \right\rangle_i. \tag{25}$$

We can therefore define the quantity $q := \left\langle \hat{\phi}'(\hat{h}_i) \right\rangle_i$ as the expectation of $\hat{\phi}'(\hat{h}_i)$ over neurons. As a consequence, $m^\mu$ can be written as

$$m^\mu = \frac{1}{\gamma(1-\gamma)(1-q)N} \sum_{j=1}^{N} (\xi_j^\mu - \gamma)\, \hat{\phi}(\hat{h}_j). \tag{26}$$

Using this equation, we can finally compute the square of $m^\nu$ for $\nu \geq 2$:

$$(m^\nu)^2 = \frac{1}{\gamma^2(1-\gamma)^2(1-q)^2N^2} \sum_{i=1}^{N} \sum_{j=1}^{N} (\xi_i^\mu - \gamma)\, (\xi_j^\mu - \gamma)\, \hat{\phi}(\hat{h}_i)\hat{\phi}(\hat{h}_j), \tag{27}$$

$$= \frac{1}{\gamma^2(1-\gamma)^2(1-q)^2N^2} \sum_{i=1}^{N} (\xi_i^\mu - \gamma)^2\, [\hat{\phi}(\hat{h}_i)]^2, \tag{28}$$

$$= \frac{p}{\gamma(1-\gamma)(1-q)^2N}. \tag{29}$$

where $p := \left\langle \hat{\phi}^2(\hat{h}_i) \right\rangle_i$. Similarly as in Eq. (22), we used that in the double sum $\sum_{i=1}^N \sum_{j=1}^N$, only the term $i = j$ survives:

$$\sum_i^N \sum_j^N \left(\xi_i^\mu - \gamma\right)\left(\xi_j^\mu - \gamma\right) = \sum_i^N \sum_j^N \left(\xi_i^\mu \xi_j^\mu - \gamma \xi_i^\mu - \gamma \xi_j^\mu + \gamma^2\right) = \tag{30}$$

$$\sum_i^N \left[ (\xi_i^\mu)^2 - 2\gamma \xi_i^\mu + \gamma^2 + \sum_{j \neq i}^N \left(\xi_i^\mu \xi_j^\mu - \gamma \xi_i^\mu - \gamma \xi_j^\mu + \gamma^2\right) \right] \tag{31}$$

$$\tag{32}$$

where the first part for $j = i$ is en expectation $\gamma(1-\gamma)$ and the second term is in expectation $\gamma^2 - 2\gamma^2 + \gamma^2 = 0$. The standard deviation of the neuron-to-neuron variability (heterogeneity), Eq. (22), is thus

$$\sigma = \sqrt{\alpha R}, \qquad R := \frac{p}{(1-q)^2}. \tag{33}$$

As a result, the input potentials, Eq. (21), can be expressed at equilibrium as

$$\hat{h}_i = \left(\xi_i^1 - \gamma\right) m^1 + \sqrt{\alpha R} Z_i + I \cdot \xi_i^1, \tag{34}$$

where $Z_i \sim N(0,1)$ are Gaussian random variables. Therefore, we find from Eq. (19) that the overlap $m^1$ at equilibrium satisfies

$$m^1 = F_1(m^1) := \frac{1}{\gamma(1-\gamma)} \left\langle \left(\xi_i^1 - \gamma\right) \hat{\phi}(\hat{h}_i) \right\rangle_i, \tag{35}$$

where $\hat{h}_i$ is given by Eq. (34). The population averages $\langle \cdot \rangle_i$ can be treated as expectations over the independent random variables $\xi_i^1$ and $Z_i$. On the one hand, $\xi_i^1$ is a Bernoulli variable such that $\xi_i^1 = 1$ with probability $P_1 = \gamma$ and $\xi_i^1 = 0$ with probability $P_0 = 1 - \gamma$. On the other hand, $Z_i$ is a standard normal random variable with probability density $p_Z(z) = \exp(-z^2/2)/\sqrt{2\pi}$. We can therefore rewrite the population average in Eq. (35) explicitly resulting in

$$F_1(m^1) = \frac{1}{\gamma(1-\gamma)} \sum_{k=0,1} P_k \left(k - \gamma\right) \int \hat{\phi}(\hat{h}_k(m^1, z)) e^{-\frac{z^2}{2}} \frac{dz}{\sqrt{2\pi}} \tag{36a}$$

where we defined

$$\hat{h}_k(m, z) = (k - \gamma)\, m + \sqrt{\alpha R(m)} z + Ik, \qquad k \in \{0, 1\}. \tag{36b}$$

$$R(m) = \frac{p(m)}{[1 - q(m)]^2} \tag{36c}$$

$$q(m) = \sum_{k=0,1} P_k \int \hat{\phi}'(\hat{h}_k(m, z)) e^{-\frac{z^2}{2}} \frac{dz}{\sqrt{2\pi}} \tag{36d}$$

$$p(m) = \sum_{k=0,1} P_k \int \hat{\phi}^2(\hat{h}_k(m, z)) e^{-\frac{z^2}{2}} \frac{dz}{\sqrt{2\pi}}. \tag{36e}$$

#### Dynamical mean-field equations

By elimating the overlaps with backround patterns in Eq. (18), the retrieval of pattern 1 can be described by a closed dynamical mean-field equation of the form

$$\hat{\tau} \frac{dm^1}{d\hat{t}} = -m^1 + F_1(m^1). \tag{37}$$

For small network load, $\alpha \ll 1$, the effect of background patterns in Eq. (34) and (36b) can be neglected. In this case, we can set $m^\nu = 0$ for $\nu \geq 2$ and it is straightforward to calculate $F_1(m^1)$. The result is Eq. (36) with $\alpha R = 0$:

$$F_1(m^1) = \frac{1}{\gamma(1-\gamma)} \sum_{k=0,1} P_k \left(k - \gamma\right) \hat{\phi}\big((k - \gamma)m + Ik\big). \tag{38}$$

For large network load $\alpha$, the effect of background patterns may not be negligible. As shown above, the equilibrium solution in this case is given by Eqs. (35) and (36). For the non-stationary dynamics Eq. (37), we still use Eq. (36) for $F_1(m^1)$ even though this equation has been derived under the assumption of stationarity. This means that we assume that the overlaps with background patters are always at their equilibrium value while the overlap variables with retrieved patterns evolve in time. While this assumption is not strictly true, it gives results in excellent agreement with full network simulations (Fig. 2C). In other words, the mean-field dynamics in Fig. 2C is correct before stimulus onset and after the system has retrieved pattern 1, whereas during transients the dynamics with $F_1(m^1)$ given by Eq. (36) is an approximation. Moreover, we argued in the discussion that assuming a small or even negligible network load $\alpha \geq 0$ is a biologically plausible assumption for the human MTL. In this case, the dynamical mean-field equations for $\alpha = 0$, Eqs. (37) and (38), are valid.

#### Mean-field equations for two correlated patterns

Taking the results derived in the previous section as a starting point, we now allow two of the stored patterns to be correlated. Without loss of generality, we choose patterns $\vec{\xi}^1$ and $\vec{\xi}^2$ to be correlated, while all other $P-2$ patterns are independent. The correlation between the two patterns is defined as the Pearson correlation coefficient (covariance/variance), in Eq. (1), which we rewrite here for convenience:

$$C = \frac{\mathrm{Cov}(\xi_i^1, \xi_i^2)}{\mathrm{Var}(\xi_i^\mu)} = \frac{P_{11} - \gamma^2}{\gamma(1-\gamma)}, \tag{39}$$

where $P_{11} = P(\xi_i^1 = 1, \xi_i^2 = 1) = \langle \xi_i^1 \cdot \xi_i^2 \rangle$ is the joint probability of a neuron to be selective to both patterns. In order to generate correlated patterns with mean activity $\langle \xi_i^1 \rangle_i = \langle \xi_i^2 \rangle_i = \gamma$ and correlation coefficient $C$, we use the indicator neuron model with $\epsilon = \Omega$ described later in the SI. However it is important to note that, if we consider only two correlated patterns, it does not matter which algorithm we choose, as it will become clear in the section "How does a network embed groups of overlapping memories?".

As in the previous section, we are interested in the retrieval dynamics of pattern $\vec{\xi}^1$. However, given the correlation with pattern $\vec{\xi}^2$, we cannot neglect the overlap of the network state with $\vec{\xi}^2$. The derivation of the system of mean-field equations in case two correlated pattern case in Eq. (41) is analogous to that described in the section above. To that end, we also assume that the stimulus only depends on the selectivities $\xi_i^1$ and $\xi_i^2$ of the neuron, i.e. $\hat{I}_i(t) = I_{\xi_i^1, \xi_i^2}(t)$ for all neurons $i = 1, \ldots, N$. The input term $h_i$ has now two non-negligible terms, both from $\vec{\xi}^1$ and $\vec{\xi}^2$:

$$\hat{h}_i(m^1, m^2) = \left(\xi_i^1 - \gamma\right) m^1 + \left(\xi_i^2 - \gamma\right) m^2 + \sqrt{\alpha R(m^1, m^2)} Z_i + I_{\xi_i^1, \xi_i^2}(t), \tag{40}$$

where

$$\hat{\tau}\frac{dm^1}{dt} = -m^1 + \frac{1}{\gamma(1-\gamma)} \left\langle \left(\xi_i^1 - \gamma\right) \hat{\phi}(\hat{h}_i) \right\rangle_i \tag{41a}$$

$$\hat{\tau}\frac{dm^2}{dt} = -m^2 + \frac{1}{\gamma(1-\gamma)} \left\langle \left(\xi_i^2 - \gamma\right) \hat{\phi}(\hat{h}_i) \right\rangle_i \tag{41b}$$

$$q = \left\langle \hat{\phi}'(\hat{h}_i) \right\rangle_i \tag{41c}$$

$$p = \left\langle \hat{\phi}^2(\hat{h}_i) \right\rangle_i \tag{41d}$$

$$R(m^1, m^2) = \frac{p}{(1-q)^2}, \tag{41e}$$

and $Z_i \sim N(0,1)$, $i = 1, \ldots, N$ are independent, standard normal random variables. Analogous to Eq. (36) we compute the population averages in Eq. (41) explicitly leading to the mean-field dynamics

$$\hat{\tau}\frac{dm^1}{dt} = -m^1 + F_1(m^1, m^2) \tag{42a}$$

$$\hat{\tau}\frac{dm^2}{dt} = -m^2 + F_2(m^1, m^2). \tag{42b}$$

Here, the nonlinear functions $F_1$ and $F_2$ are given by ($\mu = 1, 2$)

$$F_\mu(m^1, m^2) = \sum_{x^1=0,1} \sum_{x^2=0,1} \frac{x^\mu - \gamma}{\gamma(1-\gamma)} P_{x^1 x^2} \int \frac{dz}{\sqrt{2\pi}} e^{-\frac{z^2}{2}} \hat{\phi}(\hat{h}_{x^1 x^2}(m^1, m^2, z)) \tag{42c}$$

with

$$\hat{h}_{x^1x^2}(m^1,m^2,z) = \sum_{\nu=1,2} (x^\nu - \gamma)\, m^\nu + I_{x^1,x^2}(t) + \sqrt{\alpha R_h(m^1,m^2)}z. \tag{42d}$$

This function can be interpreted as the mean-field input potential of a neuron with selectivity $\xi_i^1 = x^1$ and $\xi_i^2 = x^2$, background variability $Z_i = z$, in the case when the network has overlap $m^1$ and $m^2$ with patterns 1 and 2, respectively. The last term in Eq. (42d) captures the influence of background patterns on the mean-field dynamics of $m^1(t)$ and $m^2(t)$. This influence is quantified by the function $R_h(m^1,m^2)$ representing the mean squared overlap of the system with the background patterns $\mu = 3,\ldots,P$. We used a subscript $h$ for $R_h(m^1,m^2)$ to indicate that $R$ depends functionally on the mean-field potential $\hat{h}_{x^1x^2}(m^1,m^2,z)$. This functional is given by

$$R_h(m^1,m^2) = \frac{p}{(1-q)^2} \tag{42e}$$

$$q = \sum_{x^1=0,1}\sum_{x^2=0,1} P_{x^1,x^2} \int \hat{\phi}'(\hat{h}_{x^1x^2}(m^1,m^2,z))e^{-\frac{z^2}{2}}\frac{dz}{\sqrt{2\pi}} \tag{42f}$$

$$p = \sum_{x^1=0,1}\sum_{x^2=0,1} P_{x^1,x^2} \int \hat{\phi}^2(\hat{h}_{x^1x^2}(m^1,m^2,z))e^{-\frac{z^2}{2}}\frac{dz}{\sqrt{2\pi}}. \tag{42g}$$

The mean-field input potentials $\hat{h}_{x^1x^2}(m^1,m^2,z)$, $x^1,x^2 \in \{0,1\}$, needed in Eq. (42c) are obtained from the self-consistent solution of the functional equations (42d)–(42g), details are in Section "Numerical solutions". Equations (42) simplify significantly for $\alpha = 0$, which is the parameter choice of most figures, so it is worth writing explicitly the $m^1$ and $m^2$ dynamics in the case of negligible load:

$$\begin{aligned}\hat{\tau}\frac{dm^1}{dt} =& -m^1 + \frac{1}{\gamma(1-\gamma)}\Big\{ P_{11}(1-\gamma)\hat{\phi}\left[(1-\gamma)(m^1+m^2)+I_1+I_2\right] + \\ & P_{10}(1-\gamma)\hat{\phi}\left[(1-\gamma)m^1-\gamma m^2+I_1\right] - \\ & P_{01}\gamma\hat{\phi}\left[-\gamma m^1+(1-\gamma)m^2+I_2\right] - P_{00}\gamma\hat{\phi}\left[-\gamma(m^1+m^2)\right]\Big\}, \end{aligned} \tag{43a}$$

$$\begin{aligned}\hat{\tau}\frac{dm^2}{dt} =& -m^2 + \frac{1}{\gamma(1-\gamma)}\Big\{ P_{11}(1-\gamma)\hat{\phi}\left[(1-\gamma)(m^1+m^2)+I_1+I_2\right] - \\ & P_{10}\gamma\hat{\phi}\left[-\gamma m^1+(1-\gamma)m^2+I_1\right] + \\ & P_{01}(1-\gamma)\hat{\phi}\left[(1-\gamma)m^1-\gamma m^2+I_2\right] - P_{00}\gamma\hat{\phi}\left[-\gamma(m^1+m^2)\right]\Big\}. \end{aligned} \tag{43b}$$

Here, we have used the specific form $I_{x^1,x^2}(t) = I_1(t)x^1 + I_2(t)x^2$, $x^1,x^2 \in \{0,1\}$, of the external currents, where the coefficients $I_1(t)$ and $I_2(t)$ are the external input currents given selectively to the neurons of pattern 1 and 2, respectively.

The same procedure can be generalized to generate several correlate binary patterns, as in Fig. 2B. The generalization is straightforward, we can re-write the system in Eq. (42) with one dynamical equations for each correlated pattern and add the relative terms in the input $\hat{h}(x^1,..,x^\mu,z)$. Finally we need the joint probabilities $P_{x^1,x^2,x^3}$ and $P_{x^1,x^2,x^3,x^4}$. The general formula for the joint probability is given in Eq. (102). For instance, for three correlated patterns, the mean-field dynamics analogue to Eq. (42) is given by

$$\hat{\tau}\frac{dm^1}{dt} = -m^1 + \frac{1}{\gamma(1-\gamma)}\left\langle\left(\xi_i^1-\gamma\right)\hat{\phi}(\hat{h}_i)\right\rangle_i \tag{44a}$$

$$\hat{\tau}\frac{dm^2}{dt} = -m^2 + \frac{1}{\gamma(1-\gamma)}\left\langle\left(\xi_i^2-\gamma\right)\hat{\phi}(\hat{h}_i)\right\rangle_i \tag{44b}$$

$$\hat{\tau}\frac{dm^3}{dt} = -m^3 + \frac{1}{\gamma(1-\gamma)}\left\langle\left(\xi_i^2-\gamma\right)\hat{\phi}(\hat{h}_i)\right\rangle_i \tag{44c}$$

$$q = \left\langle\hat{\phi}'(\hat{h}_i)\right\rangle_i \tag{44d}$$

$$p = \left\langle\hat{\phi}^2(\hat{h}_i)\right\rangle_i \tag{44e}$$

$$R(m^1,m^2,m^3) = \frac{p}{(1-q)^2} \tag{44f}$$

where

$$\hat{h}_i(m^1, m^2, m^3) = \left(\xi_i^1 - \gamma\right) m^1 + \left(\xi_i^2 - \gamma\right) m^2 + \left(\xi_i^3 - \gamma\right) m^3 + \sqrt{\alpha R(m^1, m^2, m^3)} Z_i + I_i. \tag{45}$$

#### Excluding self-interaction

In Section "Review: mean-field equations for independent patterns" we show the derivation of the mean-field equations for the retrieval of one pattern in an attractor neural network with self-connections ("autapses"). To make the network more biologically plausible and to avoid the creation of local minima around the attractors corresponding to the stored patterns, we now consider the case where self-interactions are excluded [8]. Indeed, not excluding the term $w_{ii} r_j$ into the input $h_i$ allows some units to self-stabilize. The effect of excluding the self-interaction term on input terms in Eqs. (36a) and (42d) is capture by a the correction term [8]:

$$\frac{q\alpha\hat{\phi}(\hat{h})}{(1-q)}. \tag{46}$$

Then, Eq. (42d) becomes

$$\hat{h}_{x^1x^2}(m^1, m^2, z) = \left(x^1 - \gamma\right) m^1 + \left(x^2 - \gamma\right) m^2 + \frac{q\alpha\phi(\hat{h}_{x^1x^2}(m^1, m^2, z))}{(1-q)} + \sqrt{\alpha r} z + I_{x^1,x^2}(t), \tag{47}$$

where again $I_{x^1,x^2}(t) = I_1(t)x^1 + I_2(t)x^2$ and $x^1, x^2 \in \{0, 1\}$. In our simulation, we used the same stimulation for both patterns, i.e. $I_1(t) = I_2(t) \equiv I(t)$. The input term in Eq. (47), is solved recursively in Fig. 2B, left hand side.

#### Correlation between background patterns

In this section, we provide the derivation of the critical correlation in the presence of correlations between background patterns (main text, Fig. 2B left).

To start with, let us suppose that each pattern is correlated with just one other, so a given pattern $\vec{\xi}^{\nu}$ is only correlated with one other pattern $\vec{\xi}^{\nu'}$, $\nu \neq \nu'$. In the following, the prime notation $\nu'$ denotes for any given pattern $\nu$ the index of the associated correlated pattern. What changes, compared to the derivation in Section "Review of attractor theory", is the variance of the heterogeneity term in Eq. (22) , $\langle \sigma^2 \rangle = \sum_{\mu,\nu>2} \left\langle \left( \xi_i^{\mu}\xi_i^{\nu} - \gamma\xi_i^{\nu} - \gamma\xi_i^{\mu} + \gamma^2 \right) \right\rangle m^{\mu} m^{\nu}$, where patterns are pair-wise correlated. For a fixed pair $(\nu, \nu')$, we obtain

$$\left\langle \left( \xi_i^{\mu}\xi_i^{\nu} - \gamma\xi_i^{\nu} - \gamma\xi_i^{\mu} + \gamma^2 \right) \right\rangle = \delta_{\mu\nu}\gamma(1-\gamma) + \delta_{\mu,\nu'}(P_{11} - \gamma^2), \tag{48}$$

$$= \gamma(1-\gamma)[\delta_{\mu,\nu} + \delta_{\mu,\nu'} C], \tag{49}$$

where the second term at the right hand side captures the effect of correlation. Background patterns can still be approximated by a Gaussian variable in the large network limit, in this case with variance:

$$\langle \sigma^2 \rangle = \gamma(1-\gamma) \sum_{\nu=3}^{P} \left[ (m^{\nu})^2 + C m^{\nu} m^{\nu'} \right]. \tag{50}$$

In order to compute Eq. (50), we need to derive $(m^{\nu})^2$ and $m^{\nu}m^{\nu'}$. In what follows, we use the same definition of $q$ and $p$ as in Eq. (36). Let us start from writing $m^{\nu}$ at the first-order Taylor expansion for $m^{\nu}$ and $m^{\nu'}$ both small:

$$F_{\nu}(m^1, \ldots, m^P) \approx \frac{1}{\gamma(1-\gamma)N} \sum_{i=1}^{N} \left(\xi_i^{\nu} - \gamma\right) \hat{\phi}(\hat{h}_i) + \frac{1}{\gamma(1-\gamma)N} \sum_{i=1}^{N} \left(\xi_i^{\nu} - \gamma\right)^2 \hat{\phi}'(\hat{h}_i) m^{\nu}$$
$$+ \frac{1}{\gamma(1-\gamma)N} \sum_{i=1}^{N} \left(\xi_i^{\nu} - \gamma\right) \left(\xi_i^{\nu'} - \gamma\right) \hat{\phi}'(\hat{h}_i) m^{\nu'} \tag{51}$$

Then, following analogous passages as Eq. (23-26) we obtain the expressions:

$$(1-q)m^{\nu} = \frac{1}{\gamma(1-\gamma)N} \sum_{i=1}^{N} \left(\xi_i^{\nu} - \gamma\right) \hat{\phi}(\hat{h}_i) + qCm^{\nu'}, \tag{52a}$$

$$(1-q)m^{\nu'} = \frac{1}{\gamma(1-\gamma)N} \sum_{i=1}^{N} \left(\xi_i^{\nu'} - \gamma\right) \hat{\phi}(\hat{h}_i) + qCm^{\nu}. \tag{52b}$$

Equation (52) is a linear system of the form:

$$Dm^{\nu} = B + qCm^{\nu'} \tag{53a}$$

$$Dm^{\nu'} = B' + qCm^{\nu} \tag{53b}$$

where $B = \frac{1}{\gamma(1-\gamma)N} \sum_{i=1}^{N} (\xi_i^{\nu} - \gamma) \hat{\phi}(\hat{h}_i)$, $D = (1-q)$, similarly, $B' = \frac{1}{\gamma(1-\gamma)N} \sum_{i=1}^{N} \left(\xi_i^{\nu'} - \gamma\right) \hat{\phi}(\hat{h}_i)$, and $C = \frac{P_{11}-\gamma^2}{\gamma(1-\gamma)}$. System Eq. (52) has solutions

$$m^{\nu} = \frac{DB + qCB'}{D^2 - C^2} \tag{54a}$$

$$m^{\nu'} = \frac{DB' + qCB}{D^2 - C^2}. \tag{54b}$$

We are now ready to write the expressions for $(m^{\nu})^2$ and $m^{\nu}m^{\nu'}$:

$$\begin{aligned} (m^{\nu})^2 =& \frac{D^2B^2 + (qC)^2(B')^2 + 2DqCBB'}{(D^2 - (qC)^2)^2}, \\ m^{\nu}m^{\nu'} =& \frac{D^2BB' + qCD(B')^2 + qCDB^2 + C^2BB'}{(D^2 - (qC)^2)^2}, \end{aligned} \tag{55}$$

where $B$ and $B'$ are analogous to the term on right hand side of Eq. (24): $(B')^2 = B^2 = \frac{p}{N\gamma(1-\gamma)}$. Note that $B^2$ and $B'^2$ are equal in expectation, however $BB' \neq B^2$ in expectation due to the correlation between $\xi^{\nu}$ and $\xi^{nu'}$. The last missing piece is the cross term $BB'$ which can also be calculated analogously to Eq. (29):

$$\begin{aligned} BB' &= \frac{1}{[N\gamma(1-\gamma)]^2} \sum_{i=1}^{N} \sum_{j=1}^{N} \left[ (\xi_i^{\nu} - \gamma) \left(\xi_j^{\nu'} - \gamma\right) \hat{\phi}(\hat{h}_i)\hat{\phi}(\hat{h}_j) \right] = \\ &= \frac{1}{[N\gamma(1-\gamma)]^2} \sum_{i=1}^{N} \left[ (P_{11} - \gamma^2) \right] \hat{\phi}^2(\hat{h}_i) = \\ &= \frac{P_{11} - \gamma^2}{N\left[\gamma(1-\gamma)\right]^2} p = \\ &= \frac{Cp}{N\gamma(1-\gamma)}. \end{aligned} \tag{56}$$

In the first passage, we used the fact that the first order approximations of $\hat{\phi}(\hat{h}_i)$ and $\hat{\phi}(\hat{h}_j)$ are independent (see Eq. (51). Plugging the expressions for $(m^{\nu})^2$ and $m^{\nu}m^{\nu'}$ into Eq. (50), we obtain the variance

$$\begin{aligned} \left\langle \sigma^2 \right\rangle =& \frac{\gamma(1-\gamma)}{(D^2 - q^2C^2)^2} \cdot \\ & \cdot \sum_{\nu \geq 3} \left\{ D^2B^2 + q^2C^2(B')^2 + 2BDqCB' + CDBB' + DqC^2B^2 + DqC^2(B')^2 + q^2C^2Bb' \right\} = \\ =& \frac{\alpha p}{(D^2 - q^2C^2)^2} \left[ D^2 + q^2C^2 + 4DqC^2 + q^2C^4 \right] \end{aligned} \tag{57}$$

Finally we can write the expression for the effective $R = \left\langle \sigma^2 \right\rangle / \alpha$, under the effect of pairwise correlation between background patterns:

$$\begin{aligned} R =& \frac{p}{\left[D^2 - (qC)\right]^2} \gamma(1-\gamma) \left[ D^2 + q^2C^2 + 4DqC^2 + q^2C^4 \right] = \\ =& \frac{p}{\left[D^2 - (qC)\right]^2} \left[ (1-q)^2 + q^2C^2 + 4(1-q)qC^2 + (1-q)^2C^4 \right]. \end{aligned} \tag{58}$$

The so obtained expression for $R$ can be substituted to that of system Eq. (41).

The derivation of Eq. (58) can be extended to the case in which background patterns share correlation $C$ between non overlapping groups of exactly $p$ patterns. To do so, we need to extend the system in Eq. (53b) linearly:

$$M = \begin{pmatrix} D & -qC & -qC & \cdots & -qC \\ -qC & D & -qC & \cdots & -qC \\ -qC & -qC & D & \cdots & -qC \\ \vdots & \vdots & \vdots & \ddots & \vdots \\ -qC & -qC & -qC & \cdots & D \end{pmatrix} \cdot \begin{pmatrix} m^{\nu} \\ m^{\nu'} \\ m_{\nu''} \\ \vdots \\ m_{\nu^{n'}} \end{pmatrix} = \begin{pmatrix} B \\ B' \\ B'' \\ \vdots \\ B^{n'} \end{pmatrix} \tag{59}$$

where $M$ is a $p \times p$ matrix. In order to find the solution $\vec{m}^{\nu} = M^{-1}\vec{B}$ of system Eq. (59) we need to invert the matrix $M$. Indeed matrices $M$ of the form

$$\{M\}_{ij} = \begin{cases} D & \text{if } i = j \\ -qC & \text{if } i \neq j \end{cases} \tag{60}$$

are invertible. In order to derive the inverted matrix we can rewrite the matrix $M$ as $M = A - qCvv^T$, where $A$ is diagonal with entries $A_{i,i} = D - qC$ and $v$ is a column vector of all ones. If $M$ and $A$ are both invertible, we can use the Sherman-Morrison formula:

$$M^{-1} = \left(A - qCvv^T\right)^{-1} = A^{-1} - \frac{-qCA^{-1}vv^TA^{-1}}{1 - qCv^TA^{-1}v}. \tag{61}$$

Since $A$ is diagonal, then $(A^{-1})_{i,i} = (A_{i,i})^{-1} = \frac{1}{D+qC}$. Then

$$\left\{M^{-1}\right\}_{ij} = \begin{cases} \frac{1}{D+qC} - \frac{1}{c(D-qC)^2} & \text{if } i = j \\ -\frac{1}{c(D-qC)^2} & \text{if } i \neq j \end{cases} \tag{62}$$

where the constant $c = -\frac{1}{qC} + n\frac{1}{D+qC}$. Terms can be re-arranged to obtain:

$$\left\{M^{-1}\right\}_{ij} = \frac{1}{Z} \begin{cases} D + qC - 1 & \text{if } i = j \\ -1 & \text{if } i \neq j \end{cases} \tag{63}$$

where

$$Z = \frac{-D + (n-1)qC}{qC(D + qC)^2}. \tag{64}$$

As a final note, we consider the case in which all patterns are equally correlated, then

$$\langle \sigma^2 \rangle = \sum_{\nu \geq 3} \sum_{\mu \geq 3} \left[ \frac{P}{N}\gamma(1-\gamma)(m^{\nu})^2 + \frac{P^2}{N}(P_{11} - \gamma^2)m^{\nu}m^{\mu} \right] \tag{65}$$

The second term in the variance diverges as $N \to \infty$ because $P = \alpha N$ unless $P_{11} = \gamma^2$. We conclude that, in the limit $N \to \infty$ and assuming that the ration between patterns and neurons is a finite constant $\alpha > 0$, it is not possible to allow a correlation $C > 0$ between all stored patterns.

#### Mean-field dynamics in the presence of adaptation and global feedback

In order to derive the mean-field equations for the model with adaptation and global feedback, we consider the simplest case, in which only two patterns are correlated ($\vec{\xi}^1$ and $\vec{\xi}^2$) while all the others are independent. Analogously to Section "Mean field equations for two correlated patterns", we can group neurons into four homogeneous populations (in the presence of background patterns, the neural populations will be slightly inhomogeneous): neurons that are selective to both patterns ($\xi_i^1 = \xi_i^2 = 1$), neurons selective to pattern 1 but not 2 ($\xi_i^1 = 1, \xi_i^2 = 0$), neurons selective to pattern 2 but not 1 ($\xi_i^1 = 0, \xi_i^2 = 1$) and neurons that are selective to neither pattern 1 or 2 ($\xi_i^1 = \xi_i^2 = 0$). The probability for a neuron to belong to population $(x^1, x^2)$, i.e. $\xi_i^1 = x^1$ and $\xi_i^2 = x^2$, is the joint probability $P_{x^1,x^2}$ in Eq. (98). Furthermore each population $(x^1, x^2)$ is characterized by a different firing threshold $\theta_{x^1,x^2}(t)$.

Analogous to the derivation of Eq. (42), we obtain the six-dimensional mean-field dynamics:

$$\hat{\tau}\frac{dm^1}{dt} = -m^1 + F_1(m^1, m^2, \{\hat{\theta}_{x^1x^2}\}), \tag{66a}$$

$$\hat{\tau}\frac{dm^2}{dt} = -m^2 + F_2(m^1, m^2, \{\hat{\theta}_{x^1x^2}\}), \tag{66b}$$

$$\hat{\tau}_\theta\frac{d\hat{\theta}_{x^1x^2}}{dt} = -\theta_{x^1x^2} + \hat{\theta}_0 + \hat{D}_\theta \hat{r}_{x^1x^2}(m^1, m^2, \hat{\theta}_{x^1x^2}), \qquad x^1, x^2 \in \{0,1\}. \tag{66c}$$

Here, we have introduced the nonlinear functions

$$F_\mu(m^1, m^2, \{\hat{\theta}_{x^1x^2}\}) = \sum_{x^1=0,1}\sum_{x^2=0,1} P_{x^1,x^2}\frac{x^\mu - \gamma}{\gamma(1-\gamma)}\hat{r}_{x^1x^2}(m^1, m^2, \hat{\theta}_{x^1x^2}), \qquad \mu = 1,2 \tag{66d}$$

$$\hat{r}_{x^1x^2}(m^1, m^2, \hat{\theta}_{x^1x^2}) = \int \hat{\phi}(\hat{h}_{x^1x^2}(m^1, m^2, \hat{\theta}_{x^1x^2}, z))e^{-\frac{z^2}{2}}\frac{dz}{\sqrt{2\pi}}, \tag{66e}$$

with the mean-field input potential

$$\hat{h}_{x^1,x^2}(m^1, m^2, \{\hat{\theta}_{x^1x^2}\}, z) = \left(x^1 - \gamma\right)m^1 + \left(x^2-\gamma\right)m^2 + \sqrt{\alpha R}z - \hat{\theta}_{x^1x^2} - \frac{\hat{J}_0(t)}{\gamma}\sum_{k_1=0,1}\sum_{k_2=0,1} P_{k_1,k_2}\hat{r}_{k_1k_2}(m^1, m^2, \hat{\theta}_{k_1k_2}). \tag{66f}$$

and the mean squared overlap of background patterns $R$ given by

$$R = \frac{p}{(1-q)^2} \tag{66g}$$

$$q = \sum_{x^1=0,1}\sum_{x^2=0,1} P_{x^1,x^2}\int \hat{\phi}'\left(\hat{h}_{x^1,x^2}(m^1, m^2, \{\hat{\theta}_{x^1x^2}\}, z)\right)e^{-\frac{z^2}{2}}\frac{dz}{\sqrt{2\pi}} \tag{66h}$$

$$p = \sum_{x^1=0,1}\sum_{x^2=0,1} P_{x^1,x^2}\int \hat{\phi}^2\left(\hat{h}_{x^1,x^2}(m^1, m^2, \{\hat{\theta}_{x^1x^2}\}, z)\right)e^{-\frac{z^2}{2}}\frac{dz}{\sqrt{2\pi}} \tag{66i}$$

In order to obtain $\hat{r}_{x^1x^2}(m^1, m^2, \hat{\theta}_{x^1x^2})$ in Eq. (66c) and (66d), Eqs. (66e) – (66i) need be solved self-consistently (for more details, see Section "Numerical Solutions").

#### Stability of the fixed points

In order to compute the stability of the fixed points in Fig. 1S of the main text, we compute the eigenvalues of the Jacobian matrix $J$ of the $m^1 - m^2$ dynamics at the point location in the $m^1 - m^2$ plane. The Jacobian matrix is symmetric and the three independent entries are computed from Eq. (41) as:

$$J_{11}(m^1, m^2) = \frac{\partial(-m^1 + F_1(m^1, m^2))}{\partial m^1} = -1 + \frac{A}{\gamma(1-\gamma)}\left\langle\left(\xi_i^1 - \gamma\right)^2 \phi'(h_i)\right\rangle_i \tag{67}$$

$$J_{12}(m^1, m^2) = \frac{\partial(-m^1 + F_1(m^1, m^2))}{\partial m^2} = J_{21}(m^1, m^2) = \frac{A}{\gamma(1-\gamma)}\left\langle\left(\xi_i^1 - \gamma\right)\left(\xi_i^2 - \gamma\right)\phi'(h_i)\right\rangle_i \tag{68}$$

$$J_{22}(m^1, m^2) = \frac{\partial(-m^2 + F_2(m^1, m^2))}{\partial m^2} = -1 + \frac{A}{\gamma(1-\gamma)}\left\langle\left(\xi_i^2 - \gamma\right)^2 \phi'(h_i))\right\rangle_i \tag{69}$$

In the numerical computation of the J, we exploited the symmetries under exchange of $m^1$ and $m^2$: $J_{22}(m^1, m^2) = J_{11}(m^2, m^1)$ and $J_{21}(m^1, m^2) = J_{12}(m^2, m^1)$ .

Analogously to the system in Eq. (41), also the Jacobian matrix can be adapted to the case of 3 or 4 correlated pattern, using the joint probabilities in Eq. (102) and the generic forms

$$J_{\mu,\mu}(m^\mu, m^\mu) = -1 + \frac{A}{\gamma(1-\gamma)} \left\langle \left(\xi_i^\mu - \gamma\right)^2 \phi'(h_i) \right\rangle_i , \tag{70}$$

$$J_{\mu,\nu}(m^\mu, m^\nu) = \frac{A}{\gamma(1-\gamma)} \left\langle \left(\xi_i^\mu - \gamma\right)\left(\xi_i^\nu - \gamma\right) \phi'(h_i) \right\rangle_i \tag{71}$$

#### The limit case $b \to \infty$: when the gain function is an Heaviside

In the limit $b \to \infty$, the gain function converges to the Heaviside step function $\phi(h) = r_{\max}\Theta(h - h_0)$ which leads to some simplifications in the explicit writing of the mean-field system Eq. (41). First of all, we can rewrite $\phi^2(h) = r_{\max}^2\Theta(h - h_0)$ and $\phi'(h) = r_{\max}\delta(h - h_0)$, where $\delta(x)$ is the Dirac delta function. In the dimension-less notation, we would then write $\phi(h) = \Theta(h - \hat{h}_0)$, $\phi^2(h) = \Theta(h - \hat{h}_0)$ and $\phi'(h) = \delta(h - \hat{h}_0)$, where $\delta(x)$ We can re-write Eq. (41) as as follows:

$$\sum_{x^1=0,1} \sum_{x^2=0,1} P_{x^1x^2} \int \frac{dz}{\sqrt{2\pi}} e^{-\frac{z^2}{2}} \Theta(\hat{h}_{x^1x^2}(m^1, m^2, z) - \hat{h}_0) \tag{72}$$

$$\hat{\tau}\frac{dm^1}{dt} = -m^1 + \sum_{x^1=0,1} \sum_{x^2=0,1} \frac{x^1-\gamma}{\gamma(1-\gamma)} P_{x^1x^2} \int \frac{dz}{\sqrt{2\pi}} e^{-\frac{z^2}{2}} \Theta(\hat{h}_{x^1x^2}(m^1, m^2, z) - \hat{h}_0) \tag{73a}$$

$$\hat{\tau}\frac{dm^2}{dt} = -m^2 + \sum_{x^1=0,1} \sum_{x^2=0,1} \frac{x^2-\gamma}{\gamma(1-\gamma)} P_{x^1x^2} \int \frac{dz}{\sqrt{2\pi}} e^{-\frac{z^2}{2}} \Theta(\hat{h}_{x^1x^2}(m^1, m^2, z) - \hat{h}_0) \tag{73b}$$

$$q = \sum_{x^1=0,1} \sum_{x^2=0,1} P_{x^1x^2} \int \frac{dz}{\sqrt{2\pi}} e^{-\frac{z^2}{2}} \delta(\hat{h}_{x^1x^2}(m^1, m^2, z) - \hat{h}_0) \tag{73c}$$

$$p = \sum_{x^1=0,1} \sum_{x^2=0,1} P_{x^1x^2} \int \frac{dz}{\sqrt{2\pi}} e^{-\frac{z^2}{2}} \Theta(\hat{h}_{x^1x^2}(m^1, m^2, z) - \hat{h}_0) \tag{73d}$$

$$R(m^1, m^2) = \frac{p}{(1-q)^2}, \tag{73e}$$

where

$$\hat{h}_i(m^1, m^2) = \left(\xi_i^1 - \gamma\right) m^1 + \left(\xi_i^2 - \gamma\right) m^2 + \sqrt{\alpha R(m^1, m^2)} Z_i + I_{\xi_i^1, \xi_i^2}(t). \tag{74}$$

In the next passage the erfc function come at hands. Erfc is defined as $\mathrm{erfc}(x) = 1 - \mathrm{erf}(x)$, where erf is the error function and we use the following identity, which follows directly from the definition:

$$\int_c^\infty \frac{e^{-\frac{x^2}{2}}}{\sqrt{2\pi}} dx = \frac{1}{2}\mathrm{erfc}\left(\frac{c}{\sqrt{2}}\right). \tag{75}$$

The identity in Eq. (75) allows to rewrite the system Eq. (73) as:

$$\tau\frac{dm^1}{dt} = -m^1 + \frac{1}{2\gamma(1-\gamma)} \sum_{x^1} \sum_{x^2} P_{x^1,x^2} \left(\xi_i^1 - \gamma\right) \mathrm{erfc}\left(\frac{h_0 - \hat{h}_i}{\sqrt{2\alpha R}}\right) \tag{76a}$$

$$\tau\frac{dm^2}{dt} = -m^2 + \frac{1}{2\gamma(1-\gamma)} \sum_{x^1} \sum_{x^2} P_{x^1,x^2} \left(\xi_i^2 - \gamma\right) \mathrm{erfc}\left(\frac{h_0 - \hat{h}_i}{\sqrt{2\alpha R}}\right) \tag{76b}$$

$$q = \sum_{x^1} \sum_{x^2} P_{x^1,x^2} \frac{1}{\sqrt{2\pi}} e^{-\frac{(\hat{h}_i)^2}{\sqrt{2\alpha R}}} \tag{76c}$$

$$p = \frac{1}{2} \sum_{x^1} \sum_{x^2} P_{x^1,x^2} \mathrm{erfc}\left(\frac{h_0 - \hat{h}_i}{\sqrt{2\alpha R}}\right) \tag{76d}$$

$$R = \frac{p}{(1-q)^2}. \tag{76e}$$

It is important to make a remark on the units of the system: if we do not use the unit-less notation, then the variable $q$ is proportional to $r_{\max}$ and the variable $p$ is proportional to $r^2_{\max}$.

If we consider the case where neural self-interaction is excluded, an extra correction term should be added to the input $h(x^1, x^2, z))$ and its limit for $b \to \infty$ reads as follows:

$$\frac{A^2 q \alpha \phi(h)}{(1 - Aq)} \xrightarrow[b\to\infty]{} \frac{A\alpha r_{\max}}{2}. \tag{77}$$

In the dimensionless notation, the correction term is reduced to a constant $\frac{\alpha}{2}$ and we can write explicitly the input term $\hat{h}_{x^1x^2}(m^1, m^2, z)$, when self interaction is excluded:

$$\hat{h}_{x^1x^2}(m^1, m^2, z) = \left(x^1 - \gamma\right) m^1 + \left(x^2 - \gamma\right) m^2 + \frac{\alpha}{2} + \sqrt{\alpha r} z + I_{x^1,x^2}(t), \tag{78}$$

Finally, in order to derive the critical correlation let us consider the retrieving state of pattern 1 (that of pattern 2 is symmetric with respect to the $m^1 - m^2$ axis in absence of external input): in this state, $m^1 = 1$, and $m^2$ depends on the correlation $C$, as it emerges from Fig. 1SA, however what is the exact value? It can be computed analytically in the limit, $b \to \infty$ and $\gamma \to 0$. We rewrite the equation for $m^2$ in Eq. 43 as:

$$\begin{aligned}\hat{\tau}\frac{dm^2}{dt} =& - m^2 + \Big\{ \frac{1}{\gamma} P_{11}(1-\gamma)\hat{\phi}\left[(1-\gamma)(m^1+m^2) + I_1 + I_2\right] - \\ & \frac{1}{1-\gamma} P_{10}\gamma\hat{\phi}\left[-\gamma m^1 + (1-\gamma)m^2 + I_1\right] + \\ & \frac{1}{\gamma} P_{01}(1-\gamma)\hat{\phi}\left[(1-\gamma)m^1 - \gamma m^2 + I_2\right] - \frac{1}{1-\gamma} P_{00}\hat{\phi}\left[-\gamma(m^1+m^2)\right]\Big\}.\end{aligned} \tag{79}$$

Next we need to write the probabilities $P_{x^1,x^2}$ as a function of $\gamma$:

$$P_{11} = \gamma^2 + \gamma(1-\gamma)C \tag{80a}$$

$$P_{10} = P_{01} = P(x^2 = 0|x^1 = 1)P(x^1 = 1) = \gamma(1-\gamma) - C\gamma(1-\gamma) \tag{80b}$$

$$P_{00} = 1 - P_{11} - P_{10} - P_{01} - P_{00} = (1-\gamma)^2 + \gamma(1-\gamma)C. \tag{80c}$$

Then, in the limit $\gamma \to 0$, we have

$$\hat{\tau}\frac{dm^2}{dt} = - m^2 + \Big\{C\hat{\phi}\left[m^1 + m^2 + I_1 + I_2\right] + (1-C)\hat{\phi}\left[m^2 + I_2\right] - \hat{\phi}\left[0\right]\Big\}. \tag{81}$$

Using the limit $b \to \infty$, in the assumption that we are recalling the first concept, $m^1 = 1$, there is not any external input, we obtain

$$m^2 = C\Theta\left(1 + m^2 - \hat{h}_0\right) + (1-C)\Theta\left(m^2 - \hat{h}_0\right) - \Theta\left(-\hat{h}_0\right). \tag{82}$$

Since $\hat{h}_0 < 1$ and $m^2 \geq 0$, the term $\Theta\left(1 + m^2 - \hat{h}_0\right) = 1$. On the other hand, $\Theta\left(-\hat{h}_0\right) = 0$. Therefore, $m^2 = C$ if $m^2 < \hat{h}_0$ (cf. the bifurcation diagram in Fig. 1SC). In the limit case where $m^2 \to \hat{h}_0$ we obtain:

$$C_{\max} \leq C_{\max} \equiv \hat{h}_0 = \frac{h_0}{A r_{\max}}. \tag{83}$$

#### Robustness to heterogeneity

##### Heterogeneous frequency-current curves

In this section we introduce heterogeneous F-I curves. Each neuron $i$ is characterised by a random baseline firing rate $r_{\min,i}$ and a random maximum firing rate $r_{\max,i}$. In the heterogeneous case, we replace the sigmoidal F-I function in Eq. 4 by

$$\phi_i(h) = \frac{r_{\max,i} - r_{\min,i}}{1 + e^{-b(h - h_{0,i})}} + r_{\min,i}. \tag{84}$$

We define two Gaussian distribution $N(\mu_{r_{\min}}, \sigma_{r_{\min}})$ and $N(\mu_{r_{\max}}, \sigma_{r_{\max}})$from which we sample $r_{\min,i}$ and $r_{\max,i}$ respectively. Since negative values of firing rates do not have a physical meaning we set $r_{\min,i}$ to

0 Hz in the case of negative values. Similarly, we do not want to allow the maximal firing rate to be too low, so we set the minimum value of the re-scaled firing rate $r_{\max,i}$ to $0.5(\mu_{r_{\max}} - \mu_{r_{\min}})$. Finally, the threshold parameter $h_{0,i}$ is defined as $h_{0,i} = h_0[r_{\max,i} - r_{\min,i}]/\mu_{r_{\max}}$, where $h_0$ is a model's parameter. Finally in Eq. 2, we re-scale the firing rates as follows:

$$r_i \to \text{Max}\left[0, \frac{r_i - r_{\min,i}}{r_{\max,i} - r_{\min,i}}\right] \mu_{r_{\max}}. \tag{85}$$

**Diluted weight matrix**

We define an attractor neural network of N units, where each unit receives input from K others. The probability of having a connection between two units is $d = M/N$. The load of the network is defined as $\alpha = P/N$, where P is the total number of patterns. We also assume that $A/d$ = constant (to be introduced into the dimensional analysis). The input term Eq. 3 is filtered with gain function $\phi$, which is chosen to be a sigmoid as in Eq. 4. The connection matrix, $w_{ij}$, contains the synaptic weights between neurons $i$ and $j$, but, compared to Eq. 5, connections are diluted with probability $d$ as defined in [1]

$$w_{ij} = \frac{A}{N\gamma(1-\gamma)} \frac{d_{ij}}{d} \sum_{\mu}^{P} (\xi_i^\mu - \gamma)\left(\xi_j^\mu - \gamma\right) \tag{86}$$

where $d_{ij}$ is 1 with probability $M/N$ and 0 otherwise and the constant $A$ can be interpret as "connection strength". In order for the weights to have expectation $\langle w_{ij}\rangle = 0$, we subtract the mean activity of patterns $< \xi_i^\mu >= \gamma$. Using the similarity measure introduced in Eq. 6, the input terms $h_j$ can also be re-written as a function of the overlaps $m^1, ..., m^P$, by using the definition of the weights $w_{ij}$, Eq. (86), and that of the overlaps.

$$\begin{aligned} h_i = \sum_j^N w_{ij} r_j &= \frac{A}{Nd\gamma(1-\gamma)} \sum_j^N d_{ij} \sum_\mu^P (\xi_i^\mu - \gamma)\left(\xi_j^\mu - \gamma\right) r_j = \\ &= \frac{A}{Nd\gamma(1-\gamma)} \sum_j^N d_{ij}\left(\xi_i^1 - \gamma\right) r_j + \frac{A}{Nd\gamma(1-\gamma)} \sum_j^N d_{ij} \sum_{\mu=2}^P \left(\xi_j^\mu - \gamma\right)(\xi_i^\mu - \gamma) r_j \end{aligned} \tag{87}$$

where we have separate the "signal" related to the first pattern being retrieved and a noise term $Y_i$. We write $h_i = Am^1 + Y_i$, with

$$Y_i = \frac{A}{Nd\gamma(1-\gamma)} \sum_j^N (1-d_{ij})\left(\xi_i^1 - \gamma\right) r_j + \frac{A}{Nd\gamma(1-\gamma)} \sum_j^N d_{ij} \sum_{\mu=2}^P \left(\xi_j^\mu - \gamma\right)(\xi_i^\mu - \gamma) r_j \tag{88}$$

Since the terms $d_{ij}$ and $\left(\xi_i^1 - \gamma\right)$ are independent, we have $\langle Y_i\rangle$.

We assume $Y_i$ to be distributed like a Gaussian with variance

$$\begin{aligned} \langle\langle \sigma^2 \rangle\rangle_i = \langle\langle (Y_i)^2 \rangle\rangle_i &= \frac{1}{N}\sum_i^N \frac{A^2}{N^2 d^2 \gamma^2 (1-\gamma)^2} \sum_{\mu\neq 1}^P \sum_{\nu\neq 1}^P \left\langle (\xi_i^\mu - \gamma)(\xi_i^\nu - \gamma) \sum_j \sum_k \left(\xi_j^\mu - \gamma\right)(\xi_k^\nu - \gamma) \right\rangle d_{ij} d_{ik} r_j r_k = \\ &= \frac{1}{N}\sum_i^N \frac{A^2}{N^2 d^2 \gamma^2 (1-\gamma)^2} \sum_{\mu\neq 1}^P (\xi_i^\mu - \gamma)^2 \sum_j d_{ij}\left(\xi_j^\mu - \gamma\right)^2 r_j^2 \end{aligned} \tag{89}$$

in the last passage, we used the fact that $\left\langle \left(\xi_j^\mu - \gamma\right)(\xi_k^\nu - \gamma)\right\rangle = \delta_{jk}\gamma(1-\gamma)$ and $d_{ij}^2 = d_{ij}$. We then apply the same independence argument as used for the signal term and obtain

$$\langle \sigma^2 \rangle = \frac{A^2 r_{\max}^2}{d^2} \gamma(1-\gamma) d \sum_\mu (m^\mu)^2 \tag{90}$$

From now on the passages are the same as in the SI, except maybe the correction term for excluding self-interaction, which I should recompute.

The final difference in the equations is that the term $\sqrt{\alpha' r} z$, where $\alpha' = P/N$ should be substituted with $\sqrt{\alpha d r} z$. The two terms however are equivalent since $\alpha' = \alpha d$.

#### How does a network embed groups of overlapping memories? Different algorithms to generate correlated patterns

In this section we describe how a single subgroup of $K$ patterns with sparseness $\gamma$ is created according to three different algorithms. Patterns belonging to the same subgroup correspond to associated concepts and share pair-wise a fraction of neurons $c$. For the hierarchical generative model and the indicator neuron model, we associate the algorithm to the theoretical probability distribution for a neuron to respond exactly to $k$ concepts out of $K$.

##### Hierarchical generative model.

We start by creating a "parent" pattern which is not part of the subgroup. The parent pattern has sparseness $\lambda = \gamma/c$: $\mathrm{prob}(\xi_i^{\mathrm{parent}} = 1) = \lambda$. We proceed to create the actual patterns by copying the ones of the parent pattern with probability $c$, while the zeros stay untouched, following the conditional probabilities

$$\mathrm{prob}(\xi_i^{\mu} = 1|\xi_i^{\mathrm{parent}} = 1) = c, \tag{91a}$$

$$\mathrm{prob}(\xi_i^{\mu} = 1|\xi_i^{\mathrm{parent}} = 0) = 1 - c, \tag{91b}$$

$$\mathrm{prob}(\xi_i^{\mu} = 1|\xi_i^{\mathrm{parent}} = 0) = 0, \tag{91c}$$

$$\mathrm{prob}(\xi_i^{\mu} = 0|\xi_i^{\mathrm{parent}} = 0) = 1. \tag{91d}$$

$$\tag{91e}$$

This ensures that the patterns $\xi_i^{\mu}$ have the right sparseness and fraction of pair-wise shared neurons. The sparseness can be checked as follows:

$$\mathrm{prob}(\xi_i^{\mu} = 1) = \lambda c = \gamma, \tag{92a}$$

$$\mathrm{prob}(\xi_i^{\mu} = 0) = \lambda(1 - c) + (1 - \lambda) = 1 - \gamma. \tag{92b}$$

On the other hand, the fraction of pair-wise shared neurons is given by the conditional probability that a neuron is part of pattern $\nu$ given that is it part of pattern $\mu$:

$$\mathrm{prob}(\xi_i^{\nu} = 1|\xi_i^{\mu} = 1) = c + (1 - c)\delta^{\mu\nu}. \tag{93}$$

Hence the fraction of shared neurons as it should be. More generically, the theoretical probability (or the expectation) that a neuron participates in $k$ patterns out of $K$ is

$$P^K(k) = \frac{K!}{(K - k)!k!}\lambda c^k (1 - c)^{K-k} + (1 - \lambda)\delta_{k0}. \tag{94}$$

##### Indicator neuron model.

To create a subgroup of pair-wise associated patterns using indicator neurons (i.e. neurons that indicate the subgroup), we proceed in three steps:

1) generate with probability $\lambda$ a small subset of indicator neurons for this subgroup. This subset gives a parent pattern of indicator neurons:

$$\mathrm{prob}(\xi_i^{\mathrm{parent}} = 1) = \lambda_{ind} = \frac{c\gamma - \gamma^2}{(1 - \epsilon)^2 - 2\gamma(1 - \epsilon) + c\gamma}. \tag{95}$$

In a network of $N$ neurons, $n_{ind} = \lambda_{ind} N$ are indicator neurons.

2) To create each pattern $\mu$ of the subgroup, copy indicator neurons with probability $(1 - \epsilon)$:

$$\mathrm{prob}(\xi_i^{\mu} = +1|\xi_i^{parent} = 1) = 1 - \epsilon \tag{96}$$

3) Add random neurons (with probability $\Omega$) to pattern $\mu$

$$\mathrm{prob}(\xi_i^{\mu} = 1|\xi_i^{\mathrm{parent}} = 0) = \Omega = \frac{\gamma - \lambda_{ind}(1 - \epsilon)}{1 - \lambda_{ind}}. \tag{97}$$

This last probability can also be interpreted as the probability of flipping a 0 from the parent pattern when creating the correlated patterns.

With this construction, the total number of neurons that are active in pattern $\mu$ is $\lambda_{ind}N(1-\epsilon) + (1-\lambda_{ind})N\frac{\gamma-\lambda_{ind}(1-\epsilon)}{1-\lambda_{ind}} = N\gamma$ as it should be. The value of $\lambda_{ind}$ is chosen in order to ensure that the fraction of pair-wise shared neurons is $c$. Indeed we found it by solving $c\gamma = \lambda(1-\epsilon)^2 + (1-\lambda)\Omega$.

In this work, we always choose $\epsilon$ such that $\epsilon = \Omega$, For specific case $\epsilon = \Omega$, it is possible to derive $\epsilon$ directly from the correlation $C$ and the sparsity $\gamma$.

We create a "parent" pattern $\vec{\xi^0}$ with mean activity $\langle \xi_i^0 \rangle_i = \lambda$. Starting from $\vec{\xi^0}$ we create $\vec{\xi^1}$ and $\vec{\xi^2}$, each unit $i$ has probability $\epsilon$ of being the equal to $\xi_i^0$ and probability $1-\epsilon$ of being flipped compared to $\xi_i^0$. All other patterns $\xi^\mu, \mu = 3, ..., P$ are sorted independently from a Bernoulli distribution with probability $P(\xi_i^\mu = 1) = \gamma$. The joint probabilities $P_{kl} = P(\xi_i^1 = k, \xi_i^2 = l)$ can be computed as functions of the probabilities $\lambda$ and $\epsilon$:

$$P_{11} = \lambda\epsilon^2 + (1-\lambda)(1-\epsilon)^2, \tag{98a}$$

$$P_{10} = P_{01} = \lambda\epsilon(1-\epsilon) + (1-\lambda)\epsilon(1-\epsilon) = \epsilon(1-\epsilon), \tag{98b}$$

$$P_{00} = \lambda(1-\epsilon)^2 + (1-\lambda)\epsilon^2. \tag{98c}$$

(98d)

Note that by this procedure we only obtain non-negative correlations $C \in [0, 1]$.

Using $P_{11}$ from Eq. (98), we can express $C$ as

$$C(\lambda, \epsilon) = \frac{P_{11} - \gamma^2}{\gamma(1-\gamma)} = \frac{(1-\lambda)\left[\lambda\epsilon^2 + (1-\epsilon)^2\right]}{\gamma(1-\gamma)}. \tag{99}$$

Similarly, the mean activity of the correlated patterns can be expressed as a function of $\lambda$ and $\epsilon$ as

$$\gamma(\lambda, \epsilon) = \langle \xi_i^1 \rangle_i = \langle \xi_i^2 \rangle_i = \lambda\epsilon + (1-\lambda)(1-\epsilon). \tag{100}$$

So far, we showed how to generate correlated patterns given the probabilities $\lambda$ and $\epsilon$. Conversely, how do we choose $\lambda$ and $\epsilon$ given the mean activity $\gamma$ and the correlation $C$, $C \geq 0$? To this end, we invert the above relations in order to solve for $\lambda(C, \gamma)$ and $\epsilon(C, \gamma)$:

$$\lambda = \frac{\gamma + \epsilon - 1}{2\epsilon - 1}, \tag{101a}$$

$$2\epsilon^3 - 3\epsilon^2 + \left[1 + 2\gamma(1-\gamma)(1-\hat{C})\right]\epsilon - \gamma(1-\gamma)(1-\hat{C}) = 0, \tag{101b}$$

Eq. (101b) has up to three solutions, we chose those that are real and in the range $[0, 1]$.

The same procedure can be generalized to generate several correlate binary patterns. The general formula for the joint probability can be written as follows:

$$P_{x^1,\ldots,x^n} = \lambda\epsilon^a(1-\epsilon)^b + (1-\lambda)\epsilon^b(1-\epsilon)^a, \tag{102}$$

where $a = \sum_{\mu=1}^n x^\mu$ is the number of $x^\mu$ variables taking value 1 and $b = n - a$ is the number of $x^\mu$ variables taking value 0. The value of the joint probabilities in Eq. (102) is invariant under permutation of the $x^\mu$.

**Iterative correlation model.**

In this subgroup construction, we do not define any parent pattern. We define the number of active neurons as $\gamma N$ and the number of pair-wise shared neurons as $\gamma c N$.

1) We define the set of "untouched neurons", which counts all neurons at the beginning of the procedure

2) We create pattern 1 by randomly sample $\gamma N$ neurons and exclude the sampled neurons from the untouched ones.

We follow the iterative steps, from 3 to 5, to create patterns 2 to p.

3) For every pattern $\xi^\mu$ with $\mu$ from 2 to $K$, compare it with each of the already created patterns. Let's suppose we are comparing the new pattern $\mu$ with the already formed pattern $\nu$. a) check how many neurons are in common between the two. b) sample from pattern $\nu$ the remaining neurons needed to reach $\gamma c N$ shared neurons.

4) Complete pattern $\mu$ by adding neurons from the untouched ones until reaching $\gamma N$ active units.

5) Remove the units used in point 4 from the untouched ones.

It is important to underline the necessity of point 3.a. To make it clearer, let's consider the case we are building a subgroup of 3 patterns. We build the first one as in point 2. When we build pattern two starting from scratch, it does not share any neuron with pattern 1, so we just sample $\gamma cN$ from pattern 1 and $\gamma N(1-c)$ from the untouched neurons. Now we move to pattern 3. As before, it does not share neurons from pattern 2, so we pick $\gamma cN$ from it. Now we compare pattern 3 with pattern 1: it ca be that between the neurons we picked from pattern 2 some belong to pattern 1 as well, that's why we need to adjust the number of neurons to pick in order to preserve the correct amount of pair-wise correlation.

When the subgroup size $K$ is big however it is still possible to exceed the correct fraction of shared neurons between some of the patterns that get build the last. Let's suppose we are creating a subgroup of size $K = 16$, I start by applying point 3 of the algorithm between pattern 16 and 15, then pattern 16 and 14 and so on. It can be that when we get to the point of picking neurons from pattern 4, 3, 2, 1 we take some neurons that also belong to pattern 15 but they are not the ones we picked in the in the previous iteration and thus get accepted. This created an higher correlation between the last built patterns in large subgroups. We checked that this does not influence significantly the average pairwise correlation during the virtual experiments described in the next section.

#### Comparing algorithm predictions with experimental data

We use a data set containing the activity of human MTL neurons [9]. Data were collected in 100 recording sessions with epileptic patients. In each recording session several stimuli were presented to the patient. The association between each pair of stimuli was estimated using a web-based association score.

In order to compare the predictions of the algorithms with the data, we try to reproduce the real data by running virtual experiments based on the three algorithms presented in the previous section. In each virtual experiment we replicate the conditions of the real experiment as follows. For each real experimental session, we first extract the number of responsive neurons in each session. We then group the presented stimuli into clusters based on an association matrix derived from the web-association scores. To do so, we use an hierarchical agglomerating clustering algorithm with threshold equal to the mean of the association matrix for the session. Such clusters indicate the amount and the size of the patterns subgroups we have to build for the corresponding virtual experiment.

We can then proceed with the virtual experiment: in each session we a) build subgroups of patterns in the same number and size as the clusters of stimuli for each of the three algorithms and then b) sample a neuron at the time and count to how many patterns does it respond to. c) Finally, the count of how many stimuli a neuron responds to that of other sessions. We sample neurons until we match the number of responsive neurons with that of the real experimental session. Each virtual experiment counts $N = 10^5$ neurons and it is run 40 times and plot in Fig. 5C the normalised mean and standard deviation.

We choose to ignore non-responding neurons in our analysis, since it is likely that the proportion of non-responsive neurons compared to that of responsive ones is largely underestimated in the experiment ( non-responsive neurons are more likely to remain silent during the experiment and not to be recorded at all).

#### Comparing virtual experiments and expected distributions

It is also possible to compare the virtual experiments with the theoretical distributions in Eq. (94) and (102). Eq. (94) and (102) provide the probability that a neuron is selective to $k$ out of $K$ patterns if a single subgroup of stimuli is stored in the network. But how do we combine such probabilities when several subgroups of patterns are stored in the network? We define $\Psi_s(k)$ the probability that a neuron responds to exactly $k$ patterns in session $s$. We know from the previous session the number and sizes $G_j$ of subgroups present in each session. Then

$$\Psi_s(k) = \sum_{j=k}^{\max K} G_j P^j(k) \zeta_j \tag{103}$$

where $\max K$ is the biggest between all subgroup sizes $K_j$ and $\zeta_j = 1 - P^j(0)$ is the probability that a neuron takes part into the subgroup $j$. The formula Eq. 103 is valid in the assumption that subgroups are strictly disjoint, meaning that we assume that the same can not take part into encoding patterns belonging to different subgroups. This assumption is not true for the way we algorithmically build subgroups patterns in the virtual experiments, however dropping it make the expression for $\Psi_s(k)$ not treatable. Finally the probabilities $\Psi_s(k)$ from each session must be combined into the final distribution $\Psi_{\text{final}}(k)$:

$$\Psi_{\text{final}}(k) = \frac{\sum_s N_s^{\text{sample}} \Psi_s(k)}{\sum_s \sum_k N_s^{\text{sample}} \Psi_s(k)} = \frac{N_1^{\text{sample}} \Psi_1(k) + N_2^{\text{sample}} \Psi_2(k) + ...}{N_{\text{tot}}^{\text{sample}}} \tag{104}$$

where $N_s^{\text{sample}}$ is the amount of responsive neurons measured in each experimental session and $N_{\text{tot}}$ is the total amount of measured responsive neurons. In the last passage, note that $\sum_s \sum_k N_s^{\text{sample}} \Psi_s(k) = \sum_s N_s^{\text{sample}} \sum_k \Psi_s(k) = N_{\text{tot}}$, since $\sum_k \Psi_s(k)$ in every session $s$. The comparison between the theoretical distributions, the virtual experiments and the experimental data is shown in Fig. 7S. The virtual experiments are the same a in Fig. 6: we re-run the experiment 40 times and took the average (main points) and standard deviation (error bars). The small mismatch between the theoretical predictions and virtual experiments is due to the fact that in the theoretical prediction we do not allow the same neurons to take part to two or more subgroups of concepts, while there is no such a restriction in the virtual experiment. Theory prediction and mean of the virtual experiments are really close, proving that only very few neurons take part in encoding different subgroups.

#### Numerical solutions

##### Two correlated patterns: finding the fixed points

The system in Eq.(41) is solved numerically to obtain the fixed nullclines, points, and flux arrows, plotted in Fig. 1S. Fixed points are obtain through a grid search in the three-dimensional space spanned by $m^1$, $m^2$ and $R$. For each value of $R_{\text{val}} \in [0, \max(R)]$ and $m^1_{\text{val}}, m^2_{\text{val}} \in [\text{Lower bound}, \text{Upper bound}]$, Eq. (42d)–(42g) are solved. We call the value of $R$ obtained by Eq. (42d) $R_{\text{reconstructed}}$. If $R_{\text{val}}$ and $R_{\text{reconstructed}}$ are close enough, namely

$$|R_{\text{val}} - R_{\text{reconstructed}}| < \text{ correction-constant} \cdot \text{step}. \tag{105}$$

The quantity called "step" is the step size of the linear space we used to span $R$,

$$\text{step} = \frac{\max(R)}{\text{Resolution}} \tag{106}$$

The correction constant can increase or decrease the range in which we accept a value $R_{\text{val}}$ as a valid solution: it is equal to 1 in most cases, but can be chose to be a bit bigger than one to avoid counting the same fixed point too many times. The values of $R_{\text{val}}$ that satisfy Eq. (105) are then used to solve Eqs. (42), providing the values $m^1_{\text{reconstructed}}$ and $m^2_{\text{reconstructed}}$. Analogously to before, we find the solutions of Eqs. (42) comparing the values $m^1_{\text{val}}$ and $m^2_{\text{val}}$ with the recomputed counterparts $m^1_{\text{reconstructed}}$ and $m^2_{\text{reconstructed}}$ as follows

$$|m^\mu_{\text{val}} - m^\mu_{\text{reconstructed}}| < \text{ correction-constant} \cdot \text{step}, \tag{107}$$

where the step is defined as

$$\text{step} = \frac{|\text{Upper bound} - \text{Lower bound }|}{\text{Resolution}}. \tag{108}$$

**List of parameters:** Fig. 1C, Fig. 1SA, Fig. 5, Fig. 4S) resolution = 1000, correction-constant = 1, size = 1000, mas(R) = 0.3, lower bound = -0.2, upper bound = 1.2. Fig. 2S) resolution = 1000, correction-constant = 1, size = 1000, mas(R) = 0.3, lower bound = -0.05, upper bound = 1.05. Fig. 2A-B) resolution = 100, correction-constant = 1.1, size = 50, mas(R) = 0.3, lower bound = -0.05, upper bound = 1.05. Fig . 3S) resolution = 500, correction-constant = 1, size = 500, mas(R) = 0.3, lower bound = -0.2, upper bound = 1.2. Fig. 6S) resolution = 500, correction-constant = 1, size = 200, mas(R) = 0, lower bound = -0.2, upper bound = 1.2.

##### Two correlated patterns with adaptation and periodic inhibition

In order to solve the dynamical equations of the mean-field in the presence of adaptation and global inhibition (as done in Fig. 4S and 5S) we compute at each point in time $\phi_{\bar{x}^1 x^2}(m^1, m^1, \Theta_{x^1 x^2})$, $p$, $q$, $R$ and $J_0(t)$. In particular, the four $\phi_{\bar{x}^1 x^2}(m^1, m^1, \Theta_{x^1 x^2})$ are solved first and recursively since they are functions of themselves. We then update $\Theta_{x^1 x^2}(t)$ with Euler method. Finally, we compute $m^1(t)$, $m^2(t)$. In order to compute $m^1(t)$, $m^2(t)$, we make a time-scale separation argument. We assume that $m^1(t)$ and $m^2(t)$ dynamics are much faster than $\Theta_{x^1x^2}(t)$ and $J_0(t)$, $\tau << T_{J_0} < T_\theta$. According to this approximation, at each point in time we let $m^1(t)$ and $m^2(t)$ reach their equilibrium values given the current $\Theta_{x^1x^2}(t)$ and $J_0(t)$. In other words, at each point in time, we consider all dynamical quantities frozen, than let $m^1(t)$ and $m^2(t)$ evolve according to their dynamics (we use Euler method) until convergence, and finally update the other quantities.

To find the fixed points in Fig. 4S, we proceed like in the non-adaptive case: we do a grid search in the space spanned by $m^1$, $m^2$ and $R$. For each solution of $R$, $\phi_{\bar{x}^1x^2}(m^1, m^1, \Theta_{x^1x^2})$ are computed recursively. Finally, for the obtained values of $R$ and $\phi_{\bar{x}^1x^2}(m^1, m^1, \Theta_{x^1x^2})$, the solutions of $m^1$ and $m^2$ are found.

##### Excluding self-interaction: a numerical approximation

In Fig. 2B, we compute the critical correlation for non-zero network load, $\alpha > 0$, in the case we consider the correction to exclude self interaction. To find the numerical solutions of the fixed points, we have approximated the input term $h(x^1, x^2, z)$ to the first order in $z$ as follows:

$$h(x^1, x^2, z) = \left\langle h(x^1, x^2, z)\right\rangle_z + A\sqrt{\alpha r}z. \tag{109}$$

Then the quantity

$$\left\langle h(x^1, x^2, z)\right\rangle_z = Ar_{\max}\left(x^1 - \gamma\right)m^1 + Ar_{\max}\left(x^2 - \gamma\right)m^2 + \left\langle \frac{A^2 q\alpha\phi(h(x^1, x^2, z))}{(1 - Aq)}\right\rangle_z \tag{110}$$

can be approximated by

$$\left\langle h(x^1, x^2, z)\right\rangle_z \sim Ar_{\max}\left(x^1 - \gamma\right)m^1 + Ar_{\max}\left(x^2 - \gamma\right)m^2 + \frac{A^2 q\alpha\phi(\left\langle h(x^1, x^2, z)\right\rangle_z)}{(1 - Aq)} \tag{111}$$

which is equivalent to take the order 0 term into the Taylor expansion of $h(x^1, x^2, z)$ for small $z$.

##### Stability of the fixed points

The stability of the fixed points in Figs. 1S,2S and 3S is obtained by computing the Jacobian matrix of the differential equations for $m^1$ and $m^2$ from Eq. (41a-b) respectively. Analogously, the stability of the fixed points in Figs. 3S, 4S are obtained by computing the Jacobian matrix of the differential equations for $m^1$ and $m^2$ from Eq. (66ca-b).

When the steepness of the gain function is very high, $b > 1000$, we approximate the gain function with an Heaviside. The system Eq. (41) as well as the Jacobian matrix are rewritten in a simpler way for $b \to \infty$ as can be found in Eq. (76).

In the numerical computation of the Jacobian matrix computed in Section "Stability of the fixed points", we exploited the symmetries under exchange of $m^1$ and $m^2$, for example $J_{22}(m^1, m^2) = J_{11}(m^2, m^1)$ and so on.

#### References


[1] Ulises Pereira and Nicolas Brunel. Attractor dynamics in networks with learning rules inferred from in vivo data. *Neuron*, 99(1):227 – 238.e4, 2018.

[2] Hugh R Wilson and Jack D Cowan. Excitatory and inhibitory interactions in localized populations of model neurons. *Biophysical journal*, 12(1):1–24, 1972.

[3] Sandro Romani, Itai Pinkoviezky, Alon Rubin, and Misha Tsodyks. Scaling laws of associative memory retrieval. *Neural computation*, 25(10):2523–2544, 2013.

[4] Stefano Recanatesi, Mikhail Katkov, Sandro Romani, and Misha Tsodyks. Neural network model of memory retrieval. *Frontiers in computational neuroscience*, 9:149, 2015.

[5] Stefano Recanatesi, Mikhail Katkov, and Misha Tsodyks. Memory states and transitions between them in attractor neural networks. *Neural computation*, 29(10):2684–2711, 2017.

[6] Michelangelo Naim, Mikhail Katkov, Sandro Romani, and Misha Tsodyks. Fundamental law of memory recall. *Physical Review Letters*, 124(1):018101, 2020.

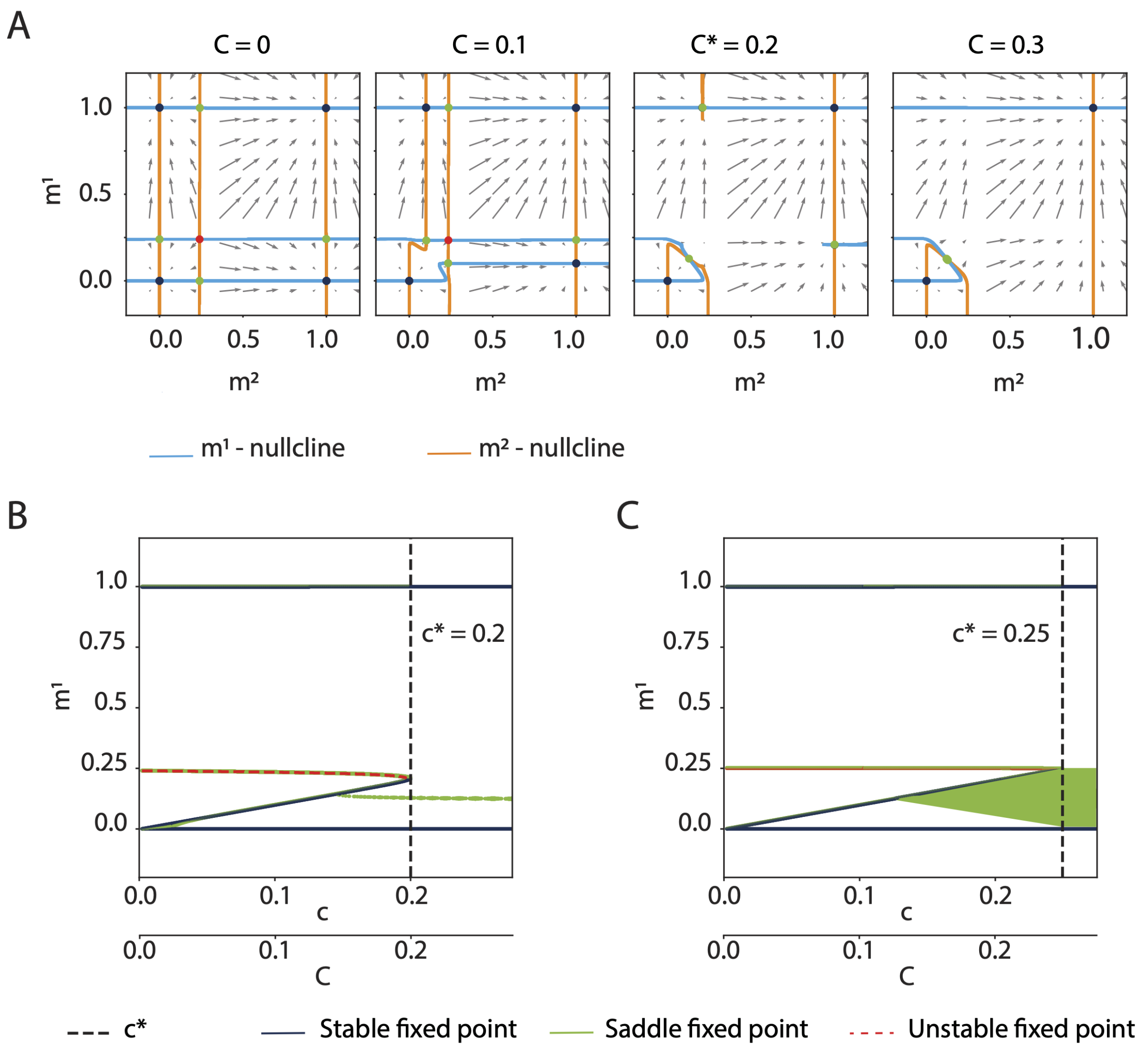


Figure 1S: A) Four phase-planes of the dynamics of variables $m^1$ and $m^2$ for different values of correlation $C$. Fixed points are color-coded by their stability: blue = stable, green = saddle and red = unstable. B) Bifurcation diagram. The projection of the fixed points position on $m^1$ is plotted against $C$. The critical correlation $C_{\max}$ is highlighted by the black dashed line. C) Same as B, but in the limit $\hat{b} \to \infty$, which leads to $C \to \hat{h}_0$. Parameters: $\gamma = 0.002$, $\hat{b} = 100$, $\hat{h}_0 = 0.25$, $\alpha = 0$.

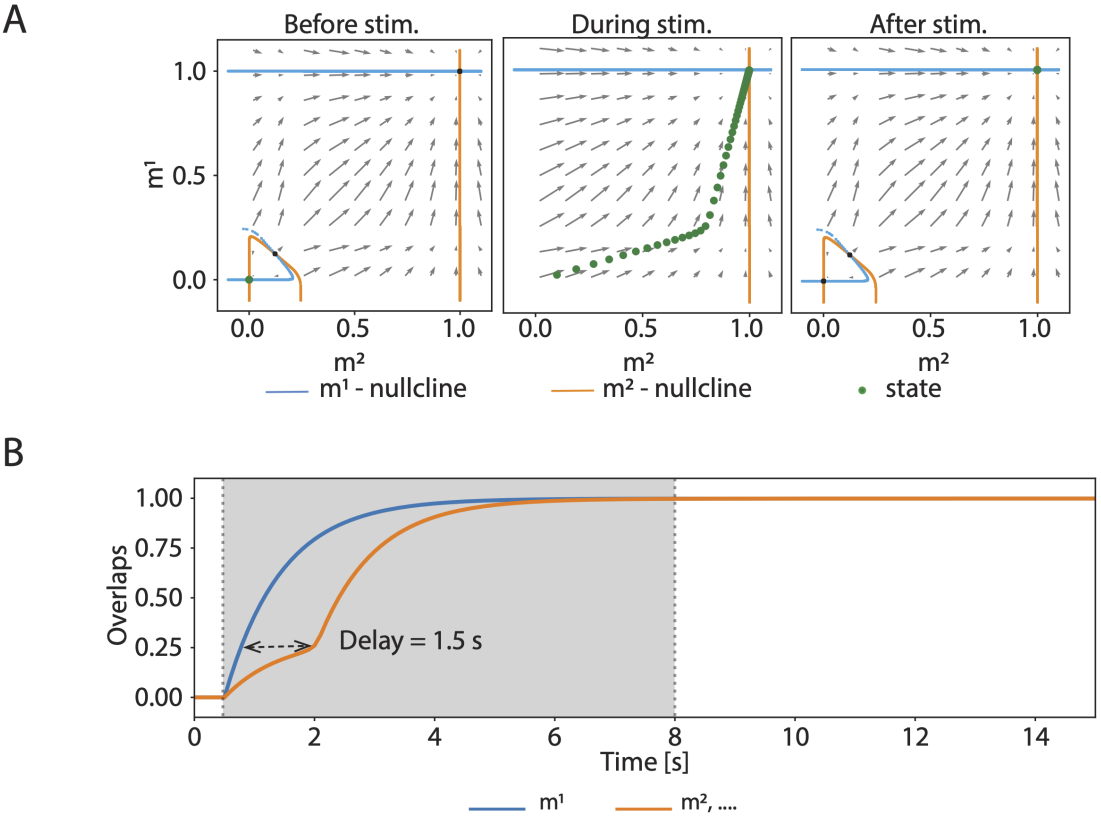


Figure 2S: Evolution of $m^1(t)$ and $m^2(t)$ according to the mean-field dynamics for super-critical correlation. The system is initialized in the rest state. During the stimulation period $(0.5-8\text{s})$ $m^1(t)$ receives external input. A) The system state is plotted in the phase-plane before, during and after stimulation respectively. B) The delay between the activation of $m^1(t)$ and $m^2(t)$ is highlighted. Parameters: $\gamma = 0.002$, $\hat{b} = 100$, $\hat{h}_0 = 0.25$, , $r_{\max} = 1$, $\hat{\tau} = \tau = 1r_{\max}$, $\alpha = 0$, $C = 0.2$.

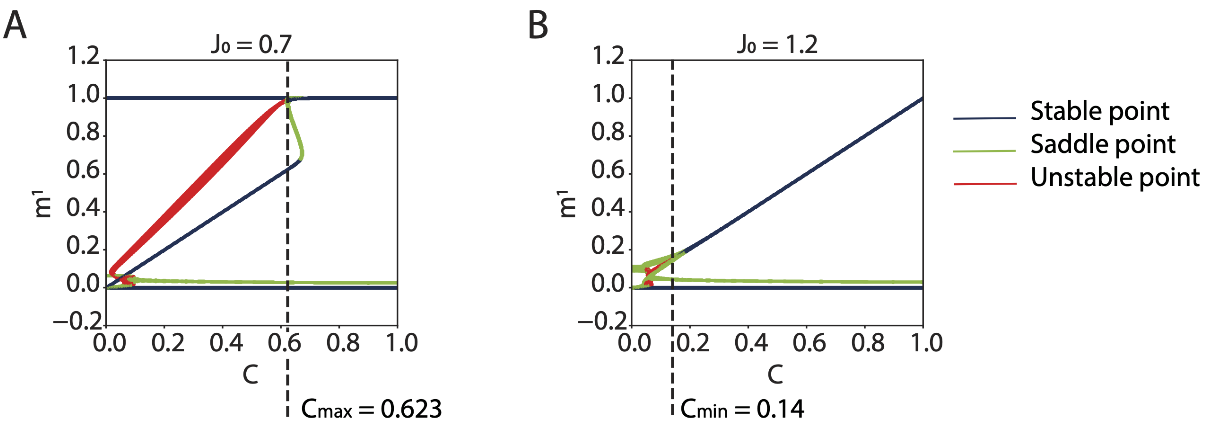


Figure 3S: Estimation of the correlation range in which retrieval of a chain of concepts is possible. A) Estimation of the maximum correlation, which correspond to the loss of the two single retrieval states, when $J_0$ is lowest. B) Estimation of the minimum correlation, which corresponds to the creation of the stable fixed point at $m^1 = m^2 > 0$, when the inhibition $J_0$ is at its maximum. In both A and B adaptation is frozen and $\theta = 0$. Parameters: $\gamma = 0.002$, $\alpha = 0$, $\hat{b} = 50$, $\hat{h}_0 = 0$, $\min(\hat{J}_0) = 0.7$, $\min(\hat{J}_0) = 1.2$.

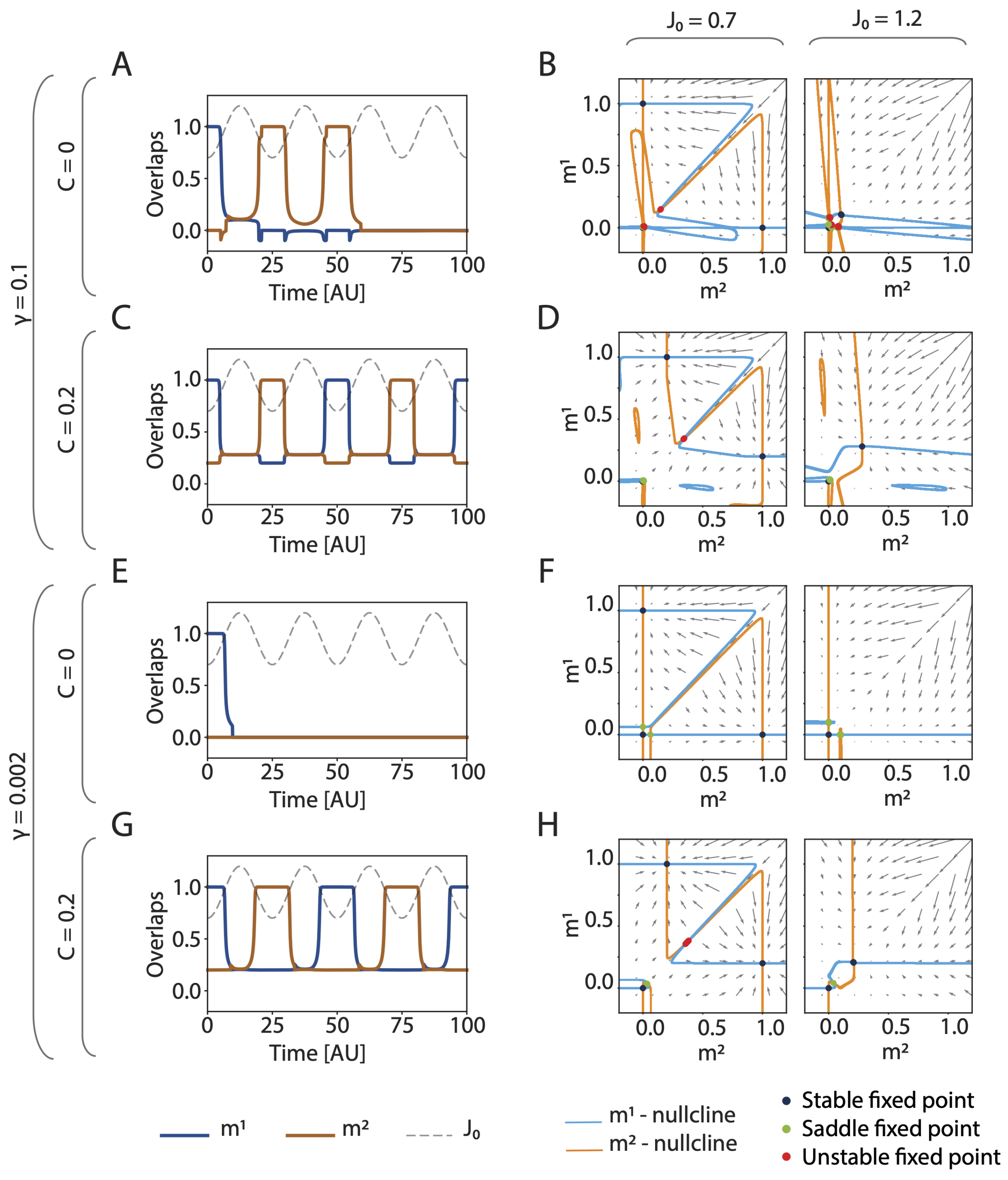


Figure 4S: A) Dynamical mean-field solutions for $m^1$ and $m^2$ in the case of two independent patterns. B) Phase planes corresponding to the minimum ($J_0 = 0.7$) and maximum ($J_0 = 1.2$) value of inhibition in the case of two independent patterns. C,D) Same as A and B, but for correlated patterns $C = 0.2$. Parameters in A - D: $\gamma = 0.1$, $\alpha = 0$, $b = 100$. E, F) Same as C and D but in the low activity regime and for independent patterns. G, H) Same as C and D but in the low activity regime. Parameters in E - H: $\gamma = 0.002$, $\alpha = 0$, $b = 100$, $\tau_\theta = 45$, $T = 0.015$, $T_{J_0} = 25$. For the dynamics: resolution = 200, factor =1. For the phase-planes: resolution = 1000, factor = 1, upper bound = 1.2, lower bound = -0.2 (same as Fig. 2 and 4.).

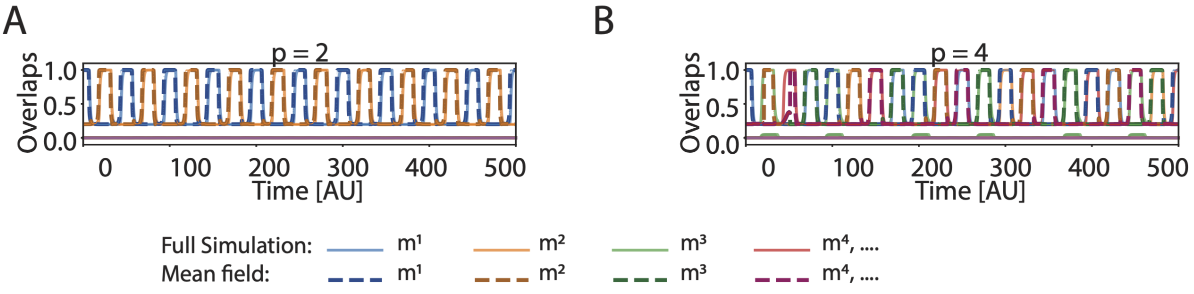


Figure 5S: Retrieval dynamics in the presence of adaptation according to the mean-field equations (dashed lines), and comparison with Fig. 4A (shaded solid lines) A) Only two patterns are correlated. B) Four patterns are correlated.Fig Parameters: $N = 10^4$, $P = 16$ in full network simulations and $\alpha = 0$ in mean-field. $\gamma = 0.002$, $\tau_\theta = 45$, $T = 0.015$, $T_{J_0} = 25$ in both.

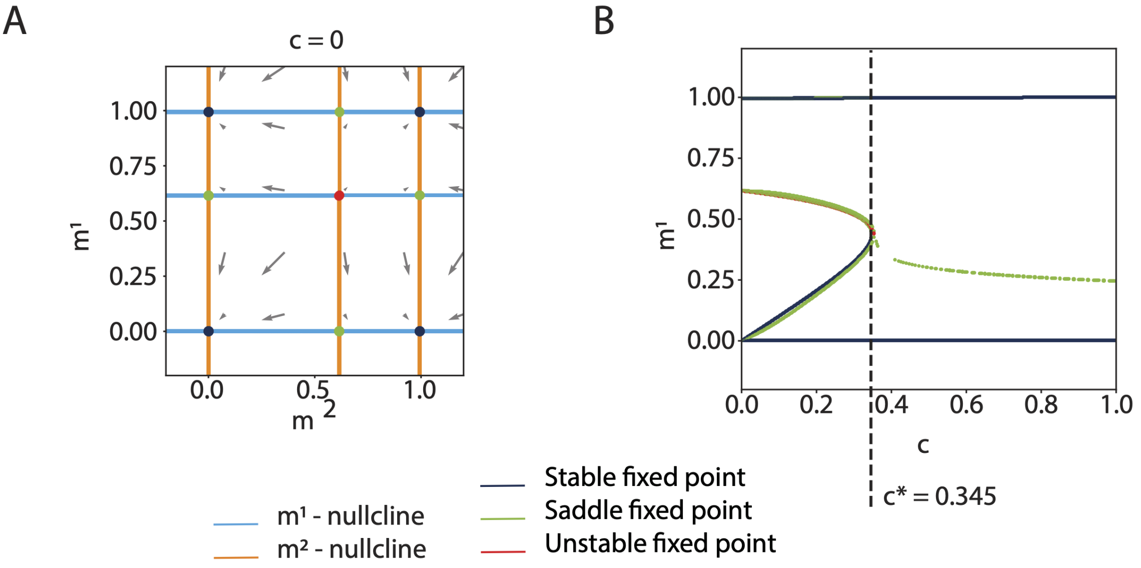


Figure 6S: Equivalent of Fig 1S but with the parameters extracted from [1]. In A and B the gain function parameters are taken as those of function $\phi$ in [1]: $A = 3.55$, $r_{\max} = 76.2$, $b = 0.82$, $h_0 = 2.46$. On the other hand, in C and D I estimated the parameters of a Sigmoid function that fits the function $f(\phi)$ in [1] as follows: $A = 3.55$, $r_{\max} = 0.83$, $b = 4.35$, $h_0 = 1.7$. In all plots $\gamma = 0.001$. A and C) The phase-plane for $c = 0$ shows the position of fixed points. B and D) Bifurcation diagram and critical fraction of shared neurons according to different parameter choices.

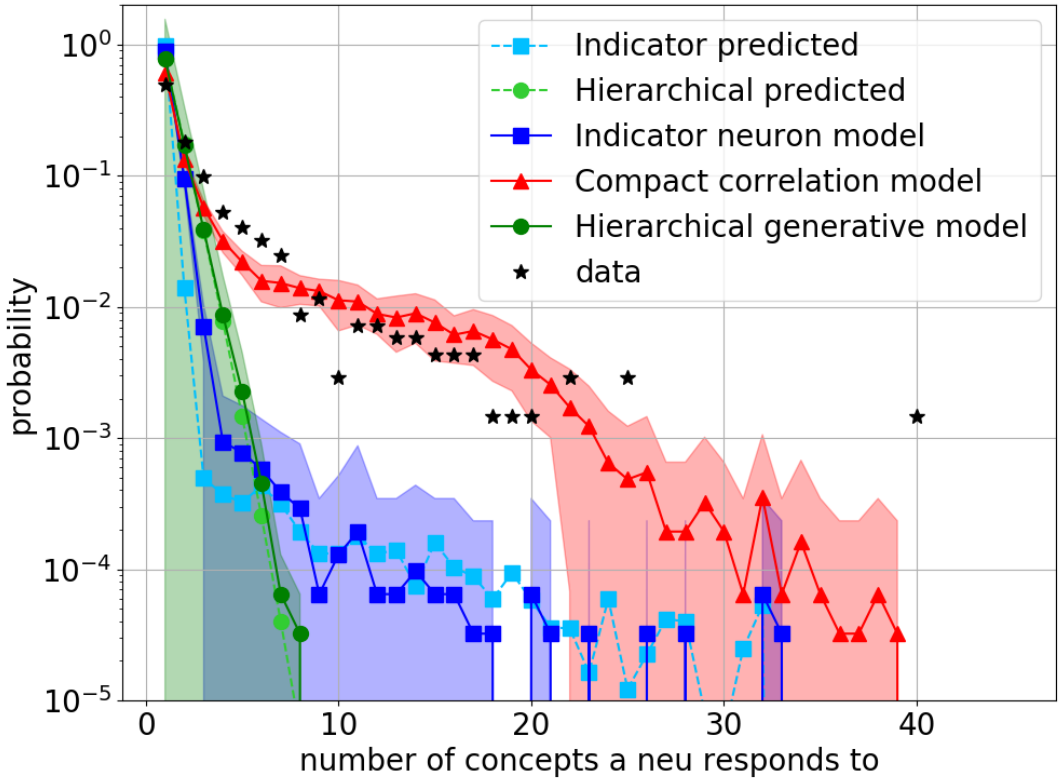


Figure 7S: Comparison between model prediction and data. Probability of finding a neuron responding to a given number of concepts as measured from experimental data (black stars), predicted by the three algorithms (as in Fig. 6, the area between error bar of one standard deviation is shaded) and theoretically forecasted for the indicator neuron model (light blue) and for the hierarchical generative model (light green) obtained from Eq. (104).

[7] Mikhail V Tsodyks and Mikhail V Feigel'man. The enhanced storage capacity in neural networks with low activity level. *EPL (Europhysics Letters)*, 6(2):101, 1988.

[8] Masatoshi Shiino and Tomoki Fukai. Self-consistent signal-to-noise analysis and its application to analogue neural networks with asymmetric connections. *Journal of Physics A: Mathematical and General*, 25(7):L375, 1992.

[9] Emanuela De Falco, Matias J Ison, Itzhak Fried, and Rodrigo Quian Quiroga. Long-term coding of personal and universal associations underlying the memory web in the human brain. *Nature communications*, 7:13408, 2016.
